# Functional diversification of the cephalopod proteome by RNA-editing

**DOI:** 10.64898/2025.12.15.694494

**Authors:** Jack M. Moen, Alicia L. Richards, Einar K. Krogsaeter, Benjamin Polacco, Martin Gordon, Erica Stevenson, Rhiana Boyles, Maximilien Burq, Dejan Stepec, Bryan Crampton, Peter Cimermancic, Joshua J. C. Rosenthal, Nevan J. Krogan, Danielle L. Swaney

## Abstract

Coleoid cephalopods exhibit the highest levels of ADAR-mediated RNA editing of any known animal, yet the functional consequences of most recoding events remain largely unknown. We integrate proteomics with biochemical and cellular assays to characterize thousands of recoding events across the *Doryteuthis pealeii* proteome. Using quantitative and functional mass spectrometry, we show that RNA edit-driven recoding reshapes the cellular proteome to alter protein stability, subcellular localization, post-translational modifications, and enzymatic activity. —Recoding can regulate post-translational modifications through their creation or ablation, and this has direct effects on protein function and protein-protein interactions. Recoding of the E3 ligase MARCHF5 drives widespread changes in substrate ubiquitylation and perturbs mitochondrial homeostasis, illustrating how RNA editing can influence organelle function. These data provide the first proteome-scale view of how extensive RNA recoding diversifies protein function in coleoid cephalopods and offers a new framework for understanding how RNA-level plasticity shapes protein function and cellular physiology.

## Introduction

RNA editing by adenosine deaminases acting on RNA (ADARs) is a widespread post-transcriptional modification capable of expanding the chemical space of RNA. Functionally, ADARs convert adenosine to inosine (A to I),^1^ a mimic for guanosine during translation.^2,3^ This single chemical modification imparts ADARs with the ability to alter protein-coding sequences, splice sites, miRNA interactions, and innate immune sensing of endogenous RNA.^4,5^ In most animals, editing acts as a regulatory buffer, fine-tuning RNA function on top of DNA-encoded sequences. Consequently, only a small number of RNA editing events in mammals are known to exert physiological effects, such as Q/R recoding of the AMPA receptor subunit GluA2, I/V recoding of kV1.1, and R/Q recoding of NEIL1, among others.^6–11^ As such, edits are considered limited and highly specialized, rather than globally defining proteome function.

Coleoid cephalopods, including octopuses, squids, and cuttlefish, possess the most complex nervous systems and behaviors among invertebrates, featuring sensorimotor processing and complex learning.^12,13^ Such complex behaviors have emerged in parallel with unusual molecular adaptations, including widespread utilization of A-to-I RNA editing in protein-coding sequences. Coleoid cephalopods exhibit tens of thousands of non-synonymous editing sites per species, representing the highest density of coding-sequence RNA editing known in the animal kingdom.^14,15^ Unlike most animals, cephalopod editing is both highly abundant and enriched within particular tissues and cellular compartments.^16^ Across coleoid species, recoding sites are conserved not only in position but also in editing level, with thousands of events under positive selection.^15^ The level of RNA editing in protein-coding regions is orders of magnitude higher in cephalopods as compared to any other species.^17^ Nearly 100,000 edits have been observed in open reading frames across several cephalopod species, while only ∼1,000 are observed in Homo sapiens and Drosophila melanogaster, and ∼100 in Mus musculus.^18^ Coleoid cephalopod ADARs are not unique in their overall activity, as millions of mammalian editing events occur in transcribed non-coding regions; instead, it is their ability to target coding sequences that is unique.^17,18^ High-level ADAR editing is conserved across coleoid cephalopods and imparts an ability to recode more than half of all sense codons, providing an expansive sequence space for proteome diversification.

The scale of cephalopod RNA editing-driven recoding has led to the hypothesis that RNA editing expands the repertoire of available proteoforms, enabling fine-tuned adaptation to environmental or physiological states.^19^ Supporting this idea, acute temperature changes can rapidly modulate editing levels at thousands of sites.^20^ Yet despite these insights, the functional landscape of recoded proteins remains largely unknown. Only a small fraction of recoding events have been validated at the protein level^15^ and few have been mechanistically linked to functional outcomes, mostly in ion channels and transporters and other proteins important for nervous function outcomes.^15,19–24^ No study to date, within cephalopods or other taxa, has provided definitive evidence that the simultaneous presence of both versions of a protein confers an adaptive benefit. In fact, early work with the Q/R site in GluA2 demonstrated that hard-wiring a point mutation in the genome to mimic the edit was sufficient to restore proper function.^6^

A major open question is how extensive recoding interfaces with the broader proteomic regulatory architecture. RNA editing has often been proposed as a mechanism for rapid proteomic diversification, enabling cells to modulate protein activity and interactions without altering genomic sequences. Post-translational modifications (PTMs), such as phosphorylation, ubiquitylation, and acetylation, tune protein function through residue-specific chemical changes. RNA editing generates analogous residue substitutions, but whether these two layers interact, compete, or cooperate has not been systematically explored. Due to the size and scope of edits in coleoids, it may be that RNA edit-driven recoding displays as much heterogeneity in function as PTMs.

Technically, the field has long faced a central obstacle: although thousands of recoding sites are observed in mRNA, in most organisms, only a small subset are detectable at the protein level. In Drosophila, roughly 1,000 editing sites have been cataloged transcriptomically, but only ∼40 have been identified by mass spectrometry;^25^ similarly, early proteomic studies in squid identified just 432 recoding sites out of the ∼60,000 non-synonymous RNA edits annotated in the study.^15^ Whether this discrepancy reflects true biological regulation, proteome-level filtering, or technical limitations of discovery proteomics remains unresolved. Consequently, despite the scale of transcriptomic recoding, the functional proteomic landscape of recoding in any species remains largely uncharted.

Here, we combine functional proteomics with biochemical and cellular assays to define the proteome-wide functional consequences of ADAR-derived recoding in *Doryteuthis pealeii*. These results reveal that extensive recoding in coleoid cephalopods provides a flexible and molecular mechanism to diversify and regulate protein function, offering a new framework for understanding how RNA-level plasticity shapes cellular physiology.

## Results

### Proteomic detection of RNA editing-derived recoding sites

While ∼87k non-synonymous edits have been described at the RNA level in *D. peallei* to date,^17^ a previous effort to map such edits at the protein level confirmed only 432 sites.^15^ To address this gap in protein-level recoding site detection, we performed a deep proteomic analysis of *D. pealeii* gill and optic lobe (**Fig. 1A**). This approach utilized both peptide fractionation to improve sensitivity in the total number of proteins detected, as well as diverse proteases to improve protein sequence coverage detection (**Fig. S1A**).^26,27^ Overall, this resulted in the detection of 181,963 peptides encompassing approximately 10.5k protein groups (**Fig. 1B).**

**Figure 1.**
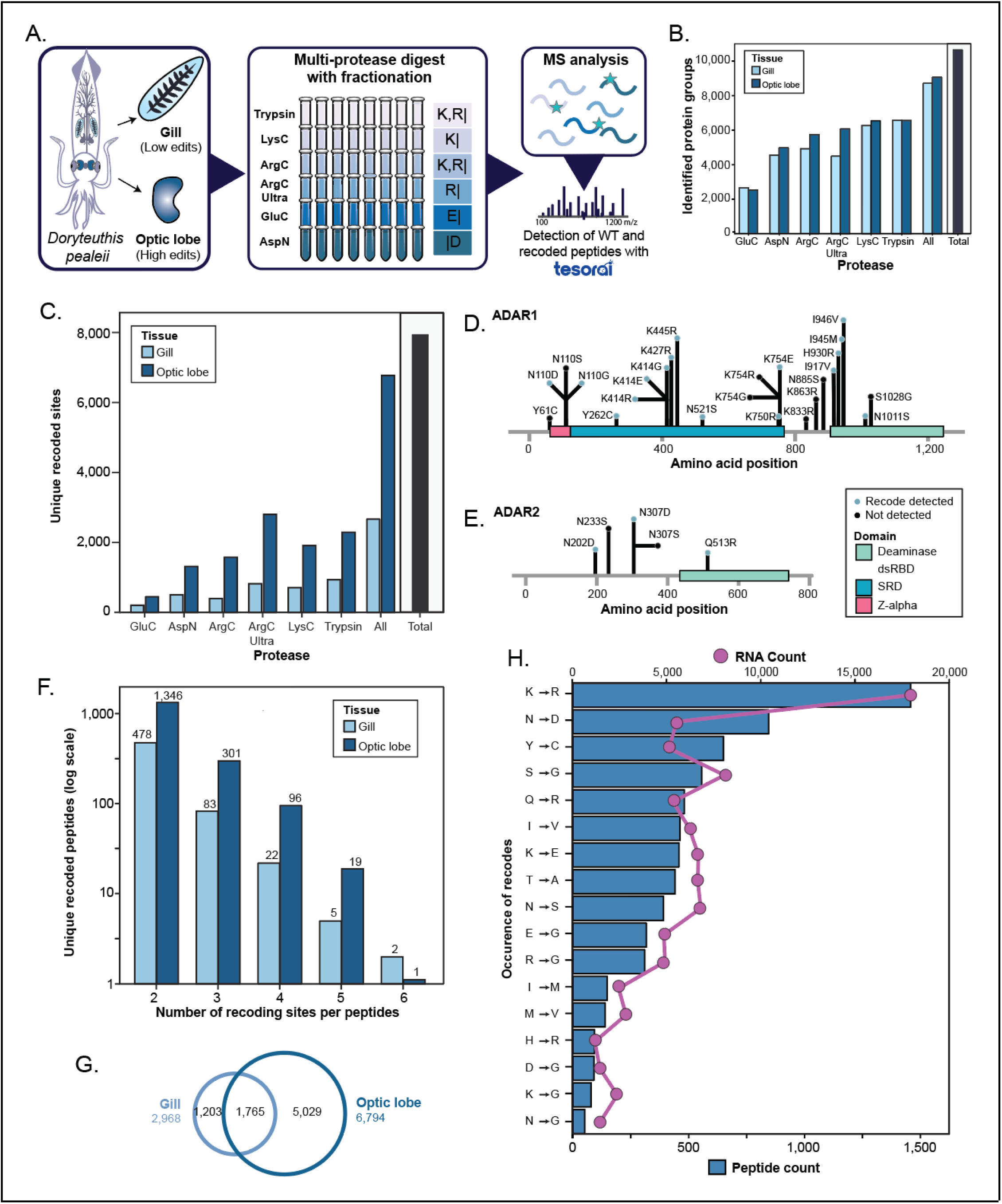
Proteomic detection of RNA editing-derived recoding sites. **A.** Workflow of the proteomic approach to detect RNA-editing induced protein recoding in the squid species of *Doryteuthis pealeii* in which peptide fractionation, diverse proteases, and the Tesorai search algorithm were utilized to enhance the detection of recoded sites. **B-C.** Number of protein groups (**B**), or recoded sites (**C**), detected across different tissue types and different proteases used for MS analysis. The total number of proteins detected across all proteases and tissue types is also displayed. **D-E.** Schematic representations of recoding sites identified for ADAR1 (**D**) and ADAR2 (**E**). **F**. Number of unique peptides identified with between 2-6 recoding sites per peptide. **G**. Overlap in protein recoding sites detected between the gill and optic lobe tissues. **H.** Ranked frequency of different recoding event types as detected by proteomics (bars) or by RNA sequencing (line).

To detect proteomic evidence of RNA-editing, we leveraged the substantial number of known edits previously detected by RNA-seq combined with Tesorai search algorithm, which employs a novel deep learning scoring approach to more robustly control false discovery rates (**Fig. S1B-C**).^28^ This resulted in the protein-level detection of 7,998 recoding sites across 2,091 proteins, representing more than an order of magnitude increase in protein-verified editing sites than previously reported (**Fig. 1C, Table S1**).^15^ This included the detection of numerous recoding events on ADAR proteins themselves, detecting 13 of 25 potential sites on ADAR1, and 3 of 5 potential sites on ADAR2 (**Fig. 1D-E**). In ADAR1, the serine-rich domain was the most heavily modified region of the protein, which is unique to coleoid cephalopods and could play a role in their high-level activity.^29^ An important advantage of our deep proteomic analysis is the ability to detect combinatorial recoding events on the same protein molecule, indicating that multiple recoding events on a given transcript are translated together. While such events have been detected through short-read RNA sequencing, it is unknown if these result in stably expressed proteins that likewise harbor multiple concurrent recoding sites. We found 1,818 peptides that contained between 2-6 recoding sites (**Fig. 1F**). Surprisingly, this included 188 proteins where the recoded sites were only observed simultaneously (i.e., both at the same time, but never alone), implying that not only are multiple sites recoded, but that there may in fact be a selection for co-occurring edits to be translated. We were also able to detect instances in which both a single and multiple recoded species occur, indicating a large pool of protein-level heterogeneity in recoded sequences.

Considering this substantial increase in recoding sites as compared to previous studies, we used several additional technical and biological metrics to benchmark our data and support the fidelity of our recoding site detections. Firstly, our use of multiple proteases provided independent detections of the same recoding site by distinct but overlapping peptide sequences, with 1,549 sites being detected by more than 1 protease, of which 23 were detected by all 6 proteases (**Fig. S1D**). Secondly, because ADAR activity is highest in the nervous system, we expect to observe a greater number of recoding sites in the optic lobe as compared to the gill.^17^ Indeed, we find that the majority of recoding sites are detected in the optic lobe (**Fig. 1G**). Lastly, RNA editing can result in a variety of different amino acid recoding events, depending upon the codon and the position of the inosine within it. Therefore, we also compared the ranked occurrence of recoding event types in both RNA and protein. Overall, we observe a highly similar trend in frequency between proteomic and RNA-seq detection, with Lys to Arg being the most common and editing of a stop codon to Trp being non-existent (**Fig. 1H**). The primary instance of discordance between editing frequencies detected by transcriptomics vs. proteomics is for Asn to Asp editing. Here, we observe an inflated frequency for Asn to Asp in our proteomics results that is likely due to a common sample preparation artifact in which Asn is converted to Asp through chemical deamidation. Based on these results, we estimate that discrepancies in Asn to Asp editing observed by proteomics may be the result of sample preparation artifacts.

### Recoding detection and prevalence

The depth of coverage from this study provides a unique opportunity to dissect factors that may influence why some edits are observed at the protein level, while others are not. Though it represents the largest detection of protein-level recoded amino acids to date, it encompasses only 9% of the 87,440 transcriptionally detected RNA editing events (**Fig. 2A**). This is perhaps not surprising, as the median level of transcriptional editing frequency across all sites is ∼1% (**Fig. S2A**), and because the proteomic abundance of peptides harboring the wild-type (WT) amino acid sequences is significantly greater than that of peptides harboring recoded sequences (**Fig. 2B**). These results indicate a general bias in the abundance for the WT amino acid across the proteome.

**Figure 2.**
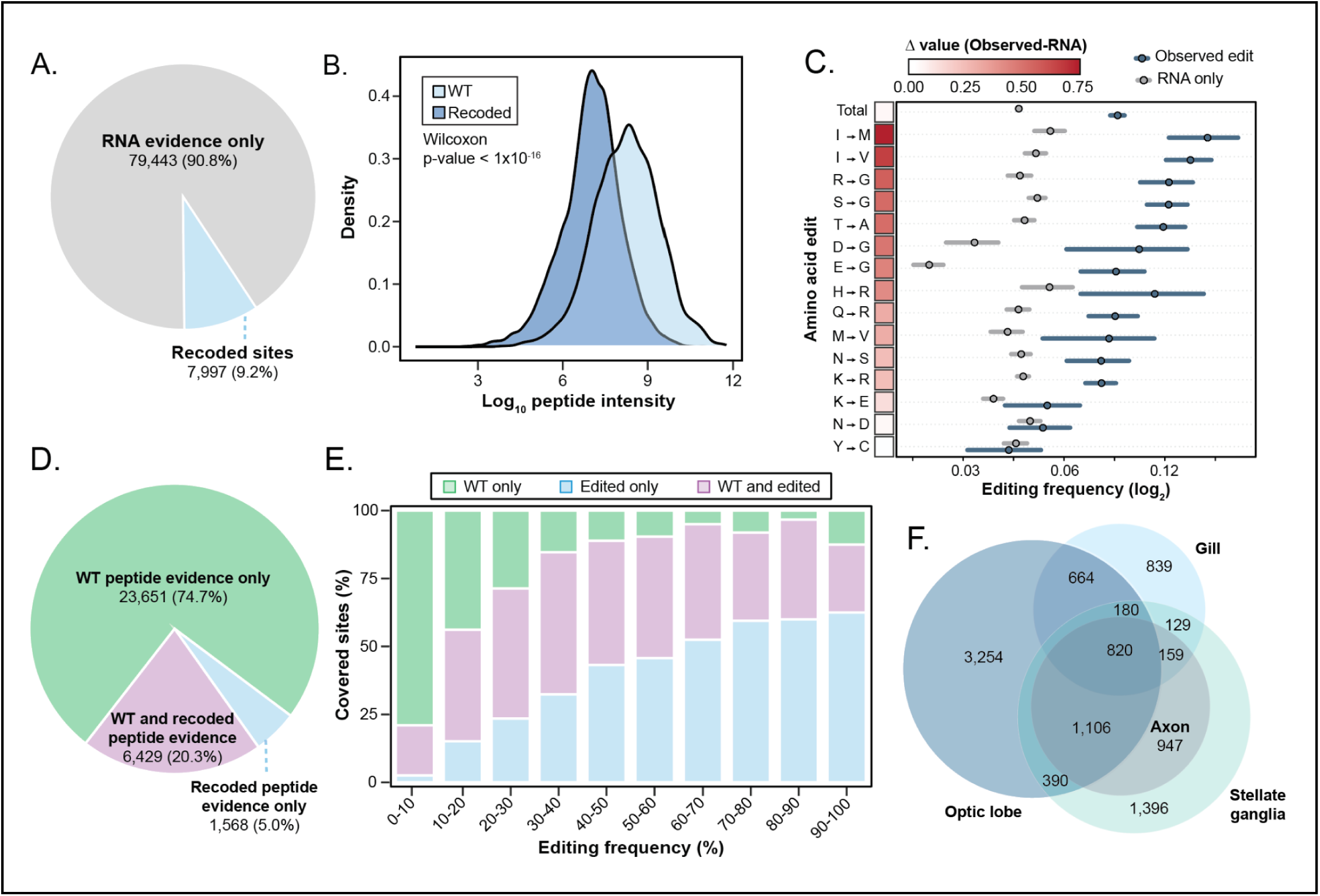
Recoding detection and prevalence. **A.** Number of possible amino acid positions that show transcript-level evidence for RNA editing and the proportion of those detected by our deep proteomics analysis. **B**. Distribution of overall peptide intensities for recoded and matching WT peptides, Wilcoxon p-value < 1x10^-16^. **C**. Comparison of RNA editing frequency observed sites (blue) against sites only seen in the RNA (grey). Heatmap shows the changes in frequency between conditions, dark outlined box indicates significance (BH corrected t-test, p-value < 0.05). **D.** Number of recoding events detected only as the WT amino acid, only as the recoded amino acid, or as being detected in both WT and recoded form. **E.** Percentage of sites covered in panel C binned by RNA editing frequency. **F.** Euler plot comparing overlapping recoded sites between the optic lobe, stellate ganglia, axons of the stellate ganglia (giant axon and surrounding small fibers), and gill.

As a whole, recoding events detected at the protein level were also biased towards more prevalent RNA editing events (**Fig. S2B**). This was true across nearly all recoding event types, with the exception of Asn to Asp and Tyr to Cys, where no such bias is observed (**Fig. 2C**). Previous studies have reported a modest correlation between peptide-level recoding and RNA editing frequency.^15^ Here we looked at all potential recoding events detected in our proteomics data and split them into 3 classes: (1) those detected only as the WT amino acid, (2) those detected in only the recoded form, and (3) those detected in both the WT and recoded forms (**Fig. 2D)**. In agreement with previous work, we also find that among editing sites with higher transcriptional frequencies, there is a shift away from the sole detection of the WT amino acid, and towards the detection of amino acids in a recoded form (**Fig. 2E**). However, it is important to note, that while this general correlation exists, we observe a high degree of individual site variability between transcriptional editing frequencies and peptide-level recoding frequencies, with numerous instances of sites being anti-correlated (**Fig. S2C**). Overall, these findings indicate that editing frequencies influence the likelihood of an edit to be detected, but does not necessarily correlate with the relative amounts seen at the protein level. This is in line with numerous proteo-genomic studies highlighting the relative discrepancies in RNAseq and protein abundance.^30^

To further probe for factors that may influence the missing coverage of edits at the protein level, we performed proteomic analysis on additional tissues. First, the stellate ganglion, a collection of soma and neuropil, and secondly, the small and giant axons that emerge from the stellate ganglia (**Fig. 2F**). As might be expected, all recoding sites detected in the axon were also detected in the stellate ganglion, indicating that substructures within a neuron may display specific editing patterns. The recoding sites detected in these two additional tissues were complementary to those previously detected in the optic lobe and gill, resulting in protein-level evidence for 2,343 additional sites. Our results indicate that some of the recoding events we were unable to observe in our initial dataset are most likely expressed in distinct neuronal subsets.

### Selective emergence of recoding events in the proteome

In order to understand the structural protein space of cephalopod proteins and their recoded proteoforms, we made use of ESM3, a protein language that prioritizes sequence, structural, and functional reasoning.^31^ In the present study, we generated structures and solvent-accessible surface area (SASA) calculations for all ∼24k proteins in the predicted proteome, as well as an additional ∼13k variants to encompass each unique combination of observed recoding sites for a given peptide. We find that amino acids that were only detected in their WT form display higher solvent accessibility compared to recoded amino acids (**Fig. 3A**). The change in SASA may be due to several factors, such as the recoding sites facilitating packing in towards the structure or changing the accessibility in a way that may increase local stability. Based on these observations, we next sought to predict the functional impact of recoded sites using AlphaMissense,^32^ which provides pathogenicity scores for all possible human amino-acid substitutions. In order to apply this framework to *D. peallei*, we aligned squid proteins to their orthologous human counterpart proteins. This analysis revealed that amino acid positions detected exclusively in the recoded form were considerably less pathogenic and consequently may display lower levels of evolutionary conservation (**Fig. 3B**). To test this hypothesis, we made use of ConSurf^33^ to generate protein-level multiple sequence alignments (MSAs) and score the evolutionary conservation of individual amino acid positions. We observe that amino acid positions detected in a recoded form have a distinct distribution, wherein the recoded amino acid is substantially less likely to be observed in alignments across species (**Fig. 3C**). These results suggest that this sub-population of recoding events is less evolutionarily conserved and when taken in conjunction with the AlphaMissense findings, implies a potential selection pressure for these events in the cephalopod proteoform space.

**Figure 3.**
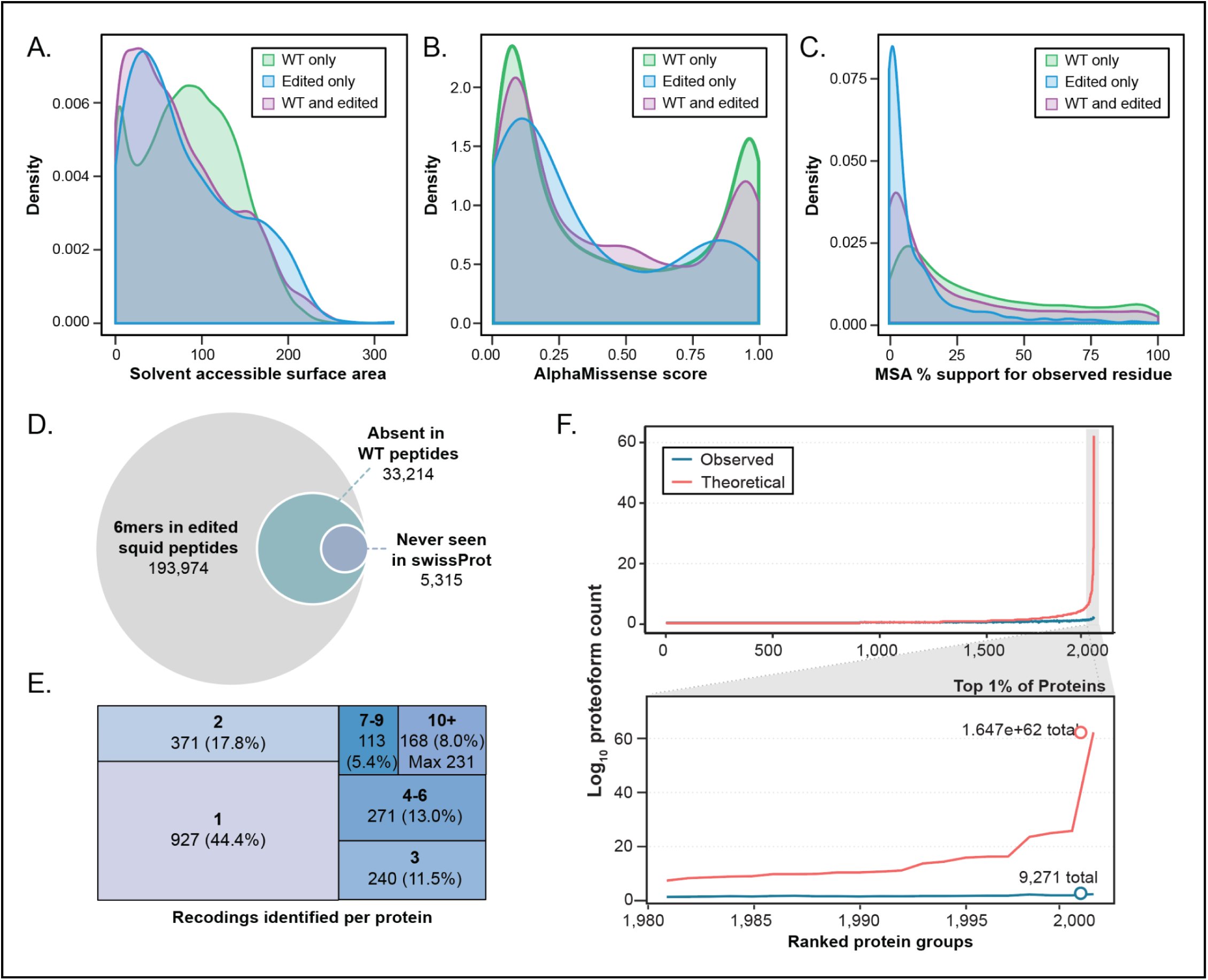
Selective emergence of recoding events in the proteome. **A.** Density plot comparing solvent accessible surface area for different classes of amino acids, as shown in Figure 2C. These include those detected only as the WT amino acid, only as the recoded amino acid, or as being detected in both WT and recoded form. **B**. Density plot for the distributions of AlphaMissense scores for *D. paellei* amino acid sites aligned to the human proteome. A higher AlphaMissense score corresponds to a higher predicted pathogenicity. Individual distributions are shown for the same amino acid classes as in panel A. **C.** Density plot for the percentage occurrence of WT or recoded residues from consurf MSAs. Individual distributions are shown for the same amino acid classes as in panel A. Amino acids were included only if they had a consurf of 0.5 or greater. For common and recoded-only amino acids, the match was selected for the resulting recoded site. **D**. Euler diagram for the number of 6 amino acid nullomers observed in edited peptide subsets. **E**. Binned groups representing the summed number of edits detected across an entire protein sequence across both gill and optic lobe tissues. **F**. Comparison of possible proteoform space from all proteins (top) or for only the top 1% most highly edited proteins (bottom).

Our confirmation of thousands of RNA editing events at the protein level has the potential to introduce entirely new amino acid sequences into the proteome. Although short amino acid sequences can theoretically exist in any sequence combination, some sequences are never observed within a given organism. These absent sequences, known as nullomers, are observed across all domains of life and are the result of unique selection pressures such as toxicity, low-abundance codons, or simply challenging sequence compositions.^34,35^ As we observe limited evolutionary conservation for numerous recoding sites, we performed a k-mer analysis with 6 amino acid windows to see if proteomically detected RNA edits may generate new amino acid sequences not present in the unmodified squid proteome. This analysis revealed that our proteomically detected recoding sites could theoretically generate 193,974 6-mers, of which 33,214 represent nullomers that are absent in the unmodified squid proteome (**Fig. 3D**). Furthermore, this extends beyond just *D. pealeii*, as 5,315 of these 6-mers are also absent from the entirety of the >14,000 species contained within the SwissProt database.^36^ These results demonstrate that RNA-editing can substantially reduce the protein-level nullomer space and enable cephalopods to express previously inaccessible protein sequences without evolving the encoding DNA.

After investigating short sequences and associated recoding events, we sought to better understand the full-length protein landscape. Looking more broadly at the frequency of recoding events across entire proteins, we were able to detect evidence of at least one recoding event on 2,091 of the possible 6,665 editable proteins we identified (**Fig. 3E**). The majority of events consisted of 1 recoding event, with a subset of proteins (168) that contained 10 or more recoding events per protein. Among the recoded proteins, we find several with a substantial number of recoding sites detected, including: spectrin alpha chain (231 sites), ankyrin-3 (102 sites), and spectrin beta chain (98 sites). The extensive number of recoding events we detected suggests that cephalopods may access one of the largest proteoform spaces of any animal. Under a conservative lower-bound model, in which each recoded peptide only produces a single recoded proteoform plus its WT counterpart, the 2,091 recoded proteins expand into 9,271 distinct proteoforms (**Fig. 3F**). Because many proteins harbor multiple recoding sites, we next evaluated the upper limit by calculating the combinatorial possibilities for each protein given the observed number of recoded positions. This resulted in an astronomical 1.6x10^62^ possible proteoforms, driven mostly by the hundreds of sites on spectrin alpha-chain. Even the second most heavily recoded protein, ankyrin-3, produced 6.2x10^25^ possible proteoforms. Given that a human cell is estimated to contain only 2x10^9^ protein molecules per cell, only a small fraction of these theoretical combinations could physically exist within a cell. Instead, these calculations reveal the scale of the proteoform space that cephalopods can theoretically explore using RNA editing and highlight the extraordinary molecular heterogeneity of this system.

### Proteoform-level analysis of recoded protein stability

The frequency of RNA editing is substantially upregulated in response to cold,^19^ and has been shown to play roles in protein stability.^19,21^ Leveraging the high-throughput nature of proteomics, we used a solvent shift assay^37^ to measure protein solubility differences between WT and recoded proteins. It is important to note that these changes in solubility can be a direct consequence of changes in protein structural stability, or can be reflective of secondary changes (e.g., changes in protein-protein interactions, or post-translational modification state). In this assay, tissue is lysed under native conditions to maintain protein structural states. The lysate is then subjected to a gradient of increasing acidic organic solvent to induce protein precipitation, and the resulting soluble proteome of each fraction is analyzed by quantitative proteomics (**Fig. 4A**). Out of 1,063 recoding sites analyzed in this assay, we found that 31% impacted protein stability and were more often destabilizing in nature (n=245) rather than stabilizing (n=87) (**Fig. 4B, Table S1**).

**Figure 4.**
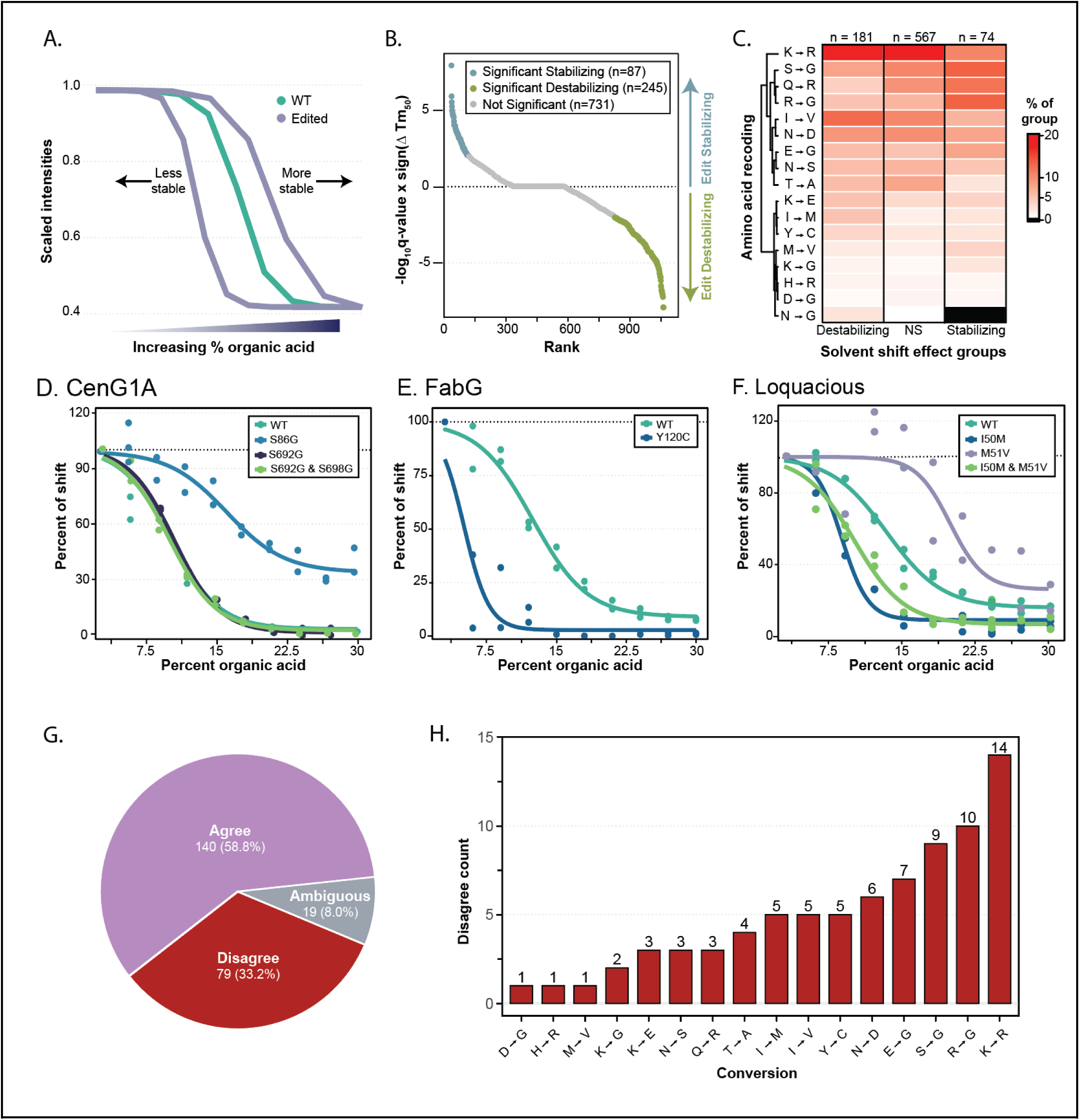
Evaluation of recoding events on protein stability. **A.** Schematic of the solvent shift method. **B.** Rank order plot for solvent shift of recoded sites ordered by the -log_10_ q-value multiplied by the direction of the shift (Recoded TM_50_ - WT TM_50_). Values with a q-value < 0.01 are colored blue (recoding is stabilizing) or green (recoding is destabilizing). **C**. Heatmap comparing the relative abundance of single recoding sites from panel B. **D-F.** Example solvent shift profiles for CenG1A (**D**), FabG (**E**), and Loquacious (**F**). Individual data points from biological replicates (n = 2) are shown for each variant along with a four-parameteric logistic regression model. **G.** Categorical comparison of observed single recoding sites between solvent shift assay and ThermoMPNN. Predicted ΔΔG and experimental ΔTm_50_ were considered in agreement if they had opposite signs with magnitudes exceeding 0.05 for both |ΔΔG| and |ΔTm_50_|. Sites with effects less than 0.05 were defined as ambiguous. **H.** Breakdown of results from panel F by recoded amino acid.

We next compared the relative amounts of each singly recoded site between those that were stabilizing, destabilizing, or non-significant (**Fig. 4C**). In line with other data, we found that Lys to Arg was the most commonly recoded site across the solvent shift data; however, this was only consistent in the destabilizing conditions, the stabilizing conditions were dominated more by Ser to Gly and Arg to Gly. When we compared the relative amounts for each group to the abundance proteomics results (**Fig. 1G**), we found that the non-significant hits were generally in agreement with the abundance proteomics IDs (**Fig. S3A**).

We find that that type of amino acid recoded does not dictate stability directly. Instead, a given recoding type (e.g., Lys to Arg) could differentially impact protein stability both across proteins as well as across amino acid positions within the same protein. As an example, we found that in CenG1A, a protein involved in synaptic transmission in *D. melanogaster,*^38^ S692G alone or in combination with S698G both display a nearly identical solubility curve to that of the WT protein (**Fig. 4D**). This may indicate that these recoding events do not impact the protein solubility of CenG1A. In contrast, the presence of the independent S698G site results in a substantial stabilization of that proteoform in comparison to WT, and may suggest that protein molecules containing this site in the absence of the nearby S692G site represent a sub-population with distinct properties. As another example, we detected a severely destabilizing Y120C site on FabG (**Fig. 4E**). This is perhaps not surprising, as Tyr to Cys recoding sites are one of the most disruptive encoded by cephalopods, with a BLOSUM62^39^ score of -2. Lastly, we detect cases for potential synergistic impacts of recoding on protein stability. For example, Loquacious harbors two adjacent recoded sites: I50M and M51V. I50M recoding resulted in decreased protein stability, while M51V substantially increased protein stability (**Fig. 4F**). Intriguingly, we also detected a peptide with concurrent recoding at both I50M and M51V. This doubly recoded event results in reduced stability as compared to WT, but it is an intermediate between either recoding site alone. This suggests that not only do individual recoding events affect protein stability, but their co-occurrence can have an has a synergistic effect.

To further assess the validity of our solvent shift data, we predicted changes in free energy for single amino acid substitutions in a given protein using ThermoMPNN.^40^ For significant single-site solvent shift recoding events, we compared whether the difference in the midpoint shift temperature (Tm_50_) aligned with the ThermoMPNN predicted change in free energy. We found that for 59% of these sites, the predicted free energy change was supportive of the solvent shift Tm_50_ change (**Fig. 4G & S3B**). This proportion of agreement is in line with the range of expected positive predictive value of the ThermoMPNN.^40^ Upon further investigation, we found that recoding to Gly produced the most discordance with ThermoMPNN; where 3 of the top 4 recoding events resulted in Gly (**Fig. 4H, S3C**). Based on the variability of Gly recoding sites observed with CenG1A (**Fig. 4C**), we believe this discrepancy is due to context-specific effects that may not be accurately predicted from the static WT protein model.

### ADAR edit-dependent recoding alters subcellular localization

To better understand the potential effects of recoding across the proteome, we first performed a GO term functional enrichment using a custom-generated STRING database for *D. peallei*.^41^ We found that for instances in which we could only detect the recoded proteoform (**Fig. 2D**), these were less likely to occur in the nucleus and more likely to be associated with the plasma membrane (**Fig. 5A**). This led us to hypothesize that ADAR edits may be able to alter subcellular localization. In order to further investigate this, we used Localization of Organelle Proteins by Isotope Tagging after Differential ultraCentrifugation (LOPIT-DC),^42^ in which a native lysate is subjected to a series of increasing centrifugation speeds to sequentially pellet varying subcellular components (**Fig. 5B**). From this, we derived 10 fractions and analyzed each fraction by mass spectrometry. Using GSEA, we were able to annotate the enrichment for various subcellular compartments across the fractions, where we observed more distinct separation of the cytosol, nucleus, and ribosome, while the plasma membrane and mitochondrial fractions were spread across several fractions (**Fig. 5C, Table S1**). We next compared the average distribution of recoded and matching WT proteins across all 10 fractions (**Fig. 5D, Fig. S4A**). Here, we observed a trend in which WT proteoforms are preferentially localized to early fractions associated with the plasma membrane and mitochondria, while recoded proteoforms are preferentially localized in the later nuclear and cytosolic fractions. In contrast, fractions 6-8 are the most similar between WT and recoded proteoforms, and encompass ribosomal proteins (fractions 7-8). This indicates that although ribosomal proteins are recoded, the recoding does not have an apparent effect on subcellular localization, presumably because they require accurate conservation and localization for function. Within each fraction, we next compared the levels of WT vs recoded proteoforms and identified 78 statistically significant differences, which primarily occurred in fraction 2 (membrane/mitochondria) and fraction 9 (nuclear) (**Fig. S4B**). As the membrane-related fractions were less clearly defined, we decided to focus on the nuclear fractions.

**Figure 5.**
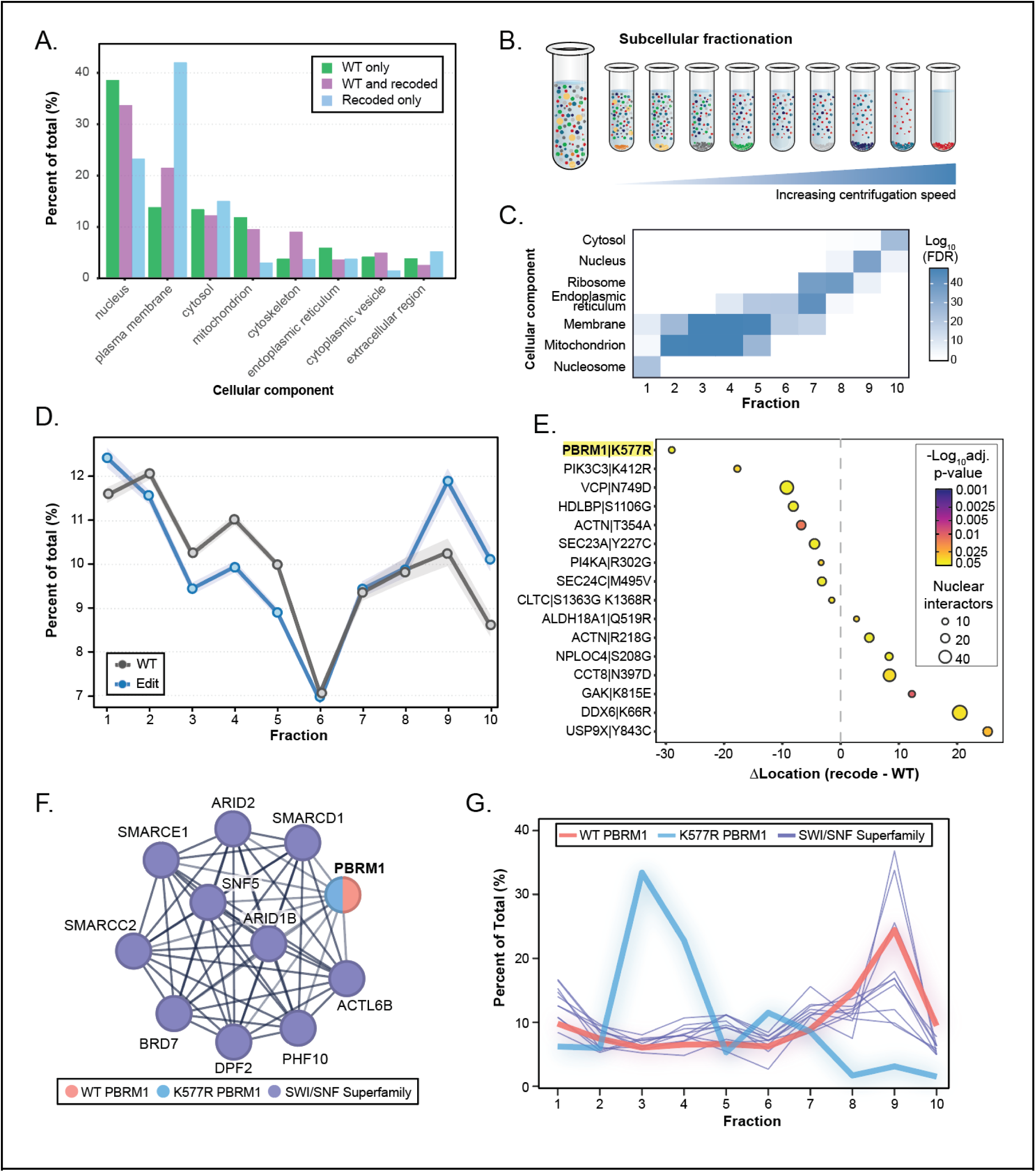
ADAR dependent recoding alters protein subcellular localization. **A.** Relative changes in subcellular localization for proteins, corresponding to observed recoding events in Fig. 2A. **B.** Schematic representation of LOPIT-DC. **C.** Heatmap of top GSEA terms by fraction. **D.** Relative percent composition of WT and corresponding recoded sequences across LOPIT fractions. Lines are denoted with a 95% confidence interval ribbon. **E.** Bubble plot comparing the change in recoded and WT composition for significant (BH corrected t-test p-value < 0.05) hits with at least ten high confidence (STRING interaction score > 600) nuclear protein interactors. Bubble size denotes the number of nuclear interactors. PBRM1 is highlighted at the top. **F.** Known PBRM1 interactors for SWI/SNF superfamily-type

Protein translocation in and out of the nucleus is a well-documented mechanism of signaling regulation; thus, we looked within fraction 9 for recoded proteoforms with a significant change in their abundance as compared to the WT protein. We further prioritized this set of proteins for those that were additionally supported by at least 10 known protein-protein interactions (PPIs) with nuclear localization (**Fig. 5E**). This narrowed down our set to 16 proteins of interest. These proteins displayed a wide variety of functions, including structural elements, post-translational modifications, and chromatin regulators. We noticed that PBRM1, a component of the PBAF complex and SWI/SNF superfamily, had the largest shift between the recoded and WT proteoforms (highlighted box **Fig. 5E**) and was positioned within a potential bipartite nuclear localization signal (NLS).^43^ When we looked at the known interactors for PBRM1, we found we were able to find WT proteins in our LOPIT that recreated a large portion of the SWI/FNF superfamily (**Fig. 5F**). The SWI/SNF complex proteins are ATP-dependent chromatin remodeling proteins that play a key role in transcription and genome stability.^44^ When we overlaid the LOPIT profiles for PBRM1 and other SWI/SNF complex members, we found they were generally in agreement, with most being highly abundant in nuclear fraction 9 (**Fig. 5G**). However, the K557R recoded proteoform of PBRM1 had almost no localization in the nucleus and was instead localized to fractions 2 and 3, the membrane/mitochondrial fractions.

### Recoded sites directly regulate post-translational modifications

Much like post-translational modifications (PTMs), RNA editing represents a dynamic modality of altering protein function. Thus, we hypothesized that RNA editing may directly influence PTM regulation by the direct regulation of modified sites, by influencing neighboring amino acids that determine substrate selectivity (i.e., motifs), or by influencing the activity and substrate selectivity of PTM-regulatory enzymes (e.g., kinases/phosphatases). Therefore, we enriched for phosphorylation, lysine acetylation, and lysine ubiquitylation modifications from both gill and optic lobe tissue (**Fig. 6A, Table S1**). From these experiments, we found a total of 23,077 unique phosphorylation sites, 4,262 lysine acetylation (AcK) sites, 13,470 lysine ubiquitylation (UbK) sites, and 65 ADP-ribosylation sites.

**Figure 6.**
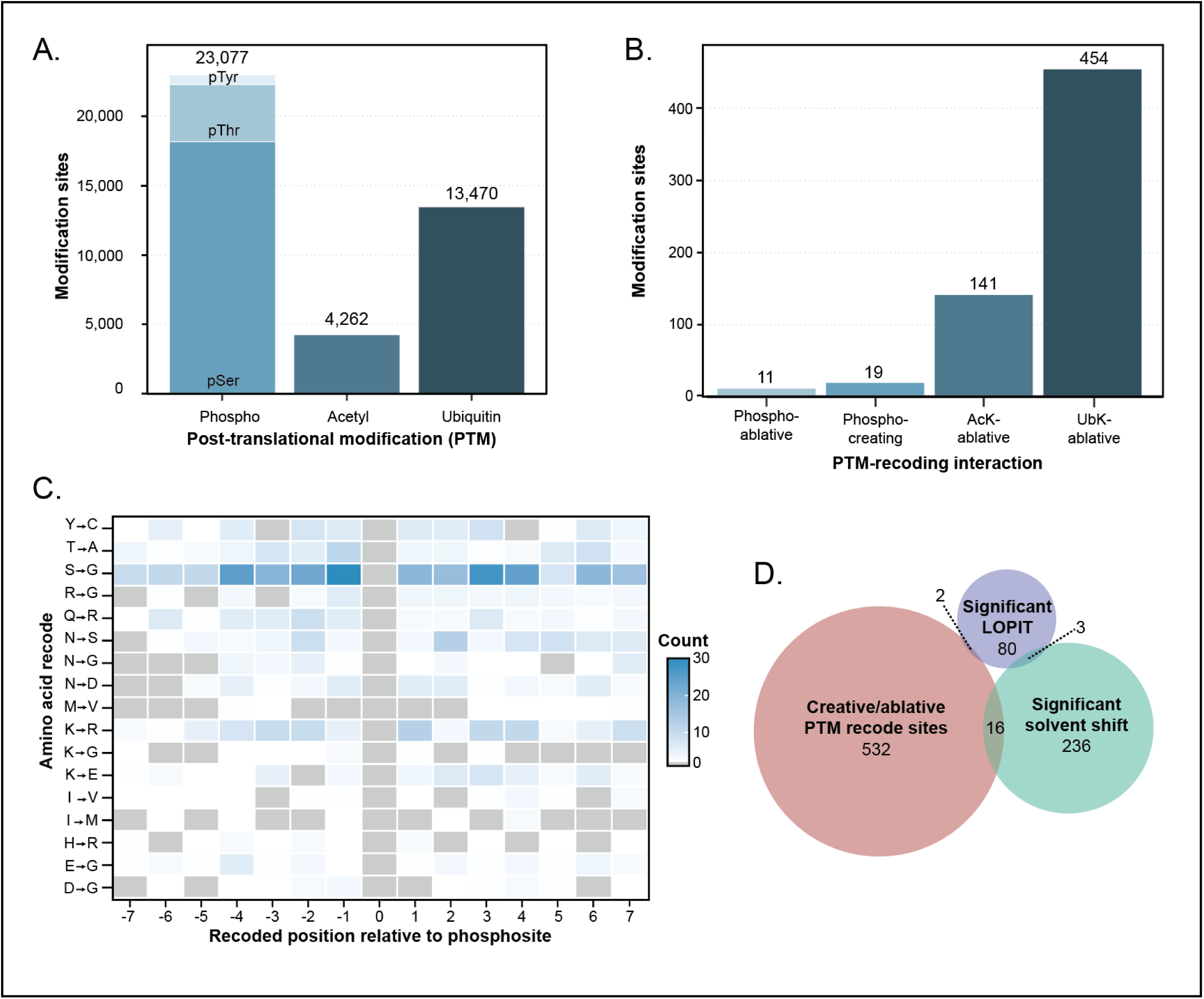
Recoding sites directly regulate post-translational modifications. **A.** Summary of PTMs identified by modification type. **B.** Direct overlap of recoding sites with PTMs broken down by interaction type. **C.** Positional scanning map of recoded only peptides in relation to the central phosphorylation site (position 0). **D.** Euler plot comparing the overlap of PTM-recoding interacting sites (red), with significant LOPIT recoding sites (purple), and significant solvent shift recoding sites (green).

One potential consequence of RNA editing is the ability to create or ablate a PTM site by recoding between different amino acids that can or cannot be modified by a given PTM. For example, Asn to Ser is a common recoding event in which the new Ser residue could be phosphorylated, while Ser to Gly recoding could prohibit phosphorylation at that position. In total, we identified 625 cases where recoding resulted in the creation or ablation of a PTM (**Fig. 6B**). These largely consisted of recoding events that ablated lysine-acetylation or lysine-ubiquitylation sites due to the high frequency of K to R edits in general (**Fig. 1G**), with only 19 instances of recoding sites that enabled the creation of a new phosphorylation site (**Fig. 6B**). This demonstrates a highly unique phenomenon associated with ADAR editing, in which a post-transcriptional modification can regulate a post-translational modification. By directly recoding such residues, cephalopods have an additional mechanism to regulate PTMs beyond their direct attachment or removal. The high number of AcK and UB ablative sites allowed us to search for amino acid motif elements using pLogo.^45^ We were only able to identify minor motif elements, such as -3 Ser for AcK sites (**Fig. S5A**) and -2 Ala for UB sites (**Fig. S5B**). This indicates that the selection for these sites is not directly driven by amino acid context, but may instead be driven by RNA or more complex protein interaction dynamics.

When looking for RNA editing-derived recoding events among these post-translationally modified peptides, there were 1,663 peptides in which a phosphorylation site was only observed in the presence of a recoding event. Such cases may indicate that, in addition to ADAR edits regulating PTM sites directly, they are also able to control the local sequence context around PTM sites (e.g., motifs), to provide an additional layer of kinase activity regulation. To further investigate this, we took this subset of phosphorylation sites and evaluated their surrounding amino acid landscape to look for recoding-dependent motif elements (**Fig. 6C**). We found Ser to Gly recoding sites were most prevalent surrounding these phosphorylation sites. Lastly, to better understand the intricacies of how the large datasets generated in the current study may compare, we looked at the overlap between significant LOPIT recoding sites, significant solvent shift recoding sites, and PTM creative/ablative sites. We observed very little overlap between the different experimental modalities, with zero recoding sites commonly detected across all three (**Fig. 6D**).

### Arginine kinase is uniquely modified in *D. peallei*

As PTM-associated recoding sites displayed the highest overlap with solvent shift sites, we sought to investigate this subset further. We noticed that one of the phospho-creative sites, N172pS, was also present in our solvent shift data in the unphosphorylated form, N172S, on arginine kinase (KARG). KARG functions as the major phosphagen regeneration system in cephalopods,^46^ creating free phospho-l-arginine as a rapid high-energy donor acting analogously to creatine kinase in humans. Considering the fact that KARG N172S can be phosphorylated, we re-analyzed our solvent shift data to additionally query for phosphorylated peptides. We observe that while the N172S recoding significantly destabilizes KARG, phosphorylation of this Ser can restore the stability to that of WT KARG (**Fig. 7A**). This suggests that a phospho-creative site may act as a compensatory modification, where phosphorylation mitigates the effect of recoding.

**Figure 7.**
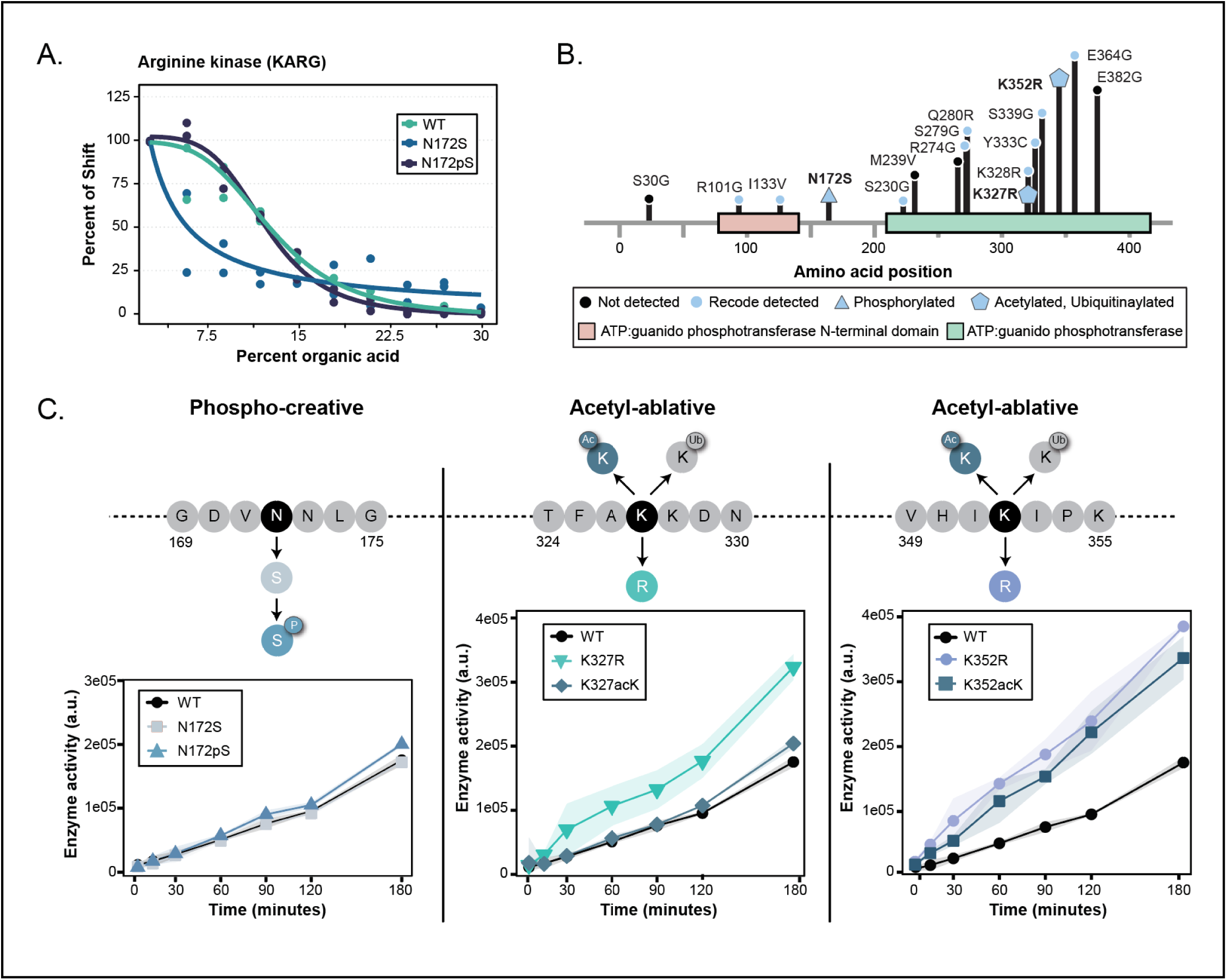
Arginine kinase is uniquely modified in D. peallei. **A.** Solvent shift profile comparing the WT, N172S recoding site, and phospho-creative N172pS recoding site. **B.** Diagram of KARG edits and PTM-recoding interacting sites detected in the study. **C.** ADP-Glo assay comparing enzymatic activity for phospho-creative (N172S, N172pS) and acetyl-ablative sites (K327R, K327acK, K352R, K352acK). With the exception of N172S, all curves were significantly different when compared to the WT, with varied effect sizes (BH corrected t-test on area under the curve, p-value < 0.05). Shaded regions surrounding the lines indicate the 95% confidence interval. Note, in some instances, the shading is not visible due to very low variance.

The highly unique effect of phosphorylation on N172S led us to further investigate the recoding and PTM landscape of KARG (**Fig. 7B**). Our proteomic analysis was able to capture a high proportion of known recoding sites on this protein (75%, 12 of 16), including two cases that were ablative for Lys acetylation or ubiquitylation (K327R, K352R). Considering that KARG displays 3 separate recoding sites that have direct interactions with PTMs and that KARG plays a central role in cephalopod biology, we theorized that the recoding events and PTMs may alter the activity of the enzyme.

To generate recombinant forms of these recoded and post-translationally modified proteins, we made use of genetic code expansion for the site-specific incorporation of phosphoserine (pS)^47^ and acetyllysine (AcK).^48^ Using the recombinant modified proteins, we performed a time course experiment assaying the generation of ADP over time (**Fig. 7C**). When comparing the same relative amounts of active kinase, we found that other than N172S, the remaining kinase variants had increased activity as compared to the WT protein, albeit with varied effect sizes. Specifically, alterations to both K327 and K352 had markedly larger impacts KARG enzymatic activity. K327R recoding enhanced activity, while acetylation of K327 (K327acK) had activity similar to the WT protein. Meanwhile, any alteration at K352 (K352R or K352acK) resulted in enhanced activity (**Fig. 7C**). In total, we observed a highly complex mixture of profiles for KARG recoding-PTM interactions, with some recoding sites and modifications having a small effect size (N172S, N172pS, K327acK) while others dramatically increased activity (K327R, K325R, K352acK). Presumably, K327 and K352 are able to alter activity as they sit in the phosphotransferase domain, while N172 sits in a linker region. Overall, this paints a picture of an impactful and complex interplay between RNA editing and PTMs that affords cephalopods an additional layer of regulation over protein function.

As we observed little to no changes in activity with the N172S, N172pS, or K327acK variants, we investigated if these alterations instead may play a role in PPIs. The same recombinant proteins as used in the activity assay were individually immobilized to beads and incubated with squid lysate to enrich for KARG PPIs. The enriched proteins were compared to the WT protein to generate a recoding-PTM specific interaction network for the low activity variants (**Fig. S6A, Table S1**). We found that K327acK had a considerably larger interaction network than the other variants. Surprisingly, the variant had decreased interactions with aerobic respiration complexes. When considered alongside the lack of changes in enzymatic activity, this implies that recoding of KARG can work in a coordinated fashion to control cellular energy production outside of directly controlling enzymatic activity. We also found that recoding the N172S site only produced nine significant interactions, two of which were recoding specific. Strikingly, N172pS interactions were more common than N172S interactions, almost exclusively upregulated compared to the WT, and some were even recoding specific interactions. Our data demonstrate that recoding N172S does not drive activity or PPIs, but that the interactions are driven by the additional phosphorylation event, highlighting a novel two-step mechanism required to facilitate PPIs. Overall, our analysis of KARG demonstrates a direct interplay between recoding and PTMs, as well as how individual sites can drive unique effects upon stability, activity, and PPIs.

### Recoding remodels organelle function

Beyond kinases, ubiquitylation signaling also encompasses a broad network of hundreds of proteins and involves a complex multi-enzyme cascade. We next looked at recoding events on different ubiquitylation machinery proteins, including E1 enzymes, E2 enzymes, and E3 ligases (**Fig. 8A**). We observed that 13 E3 ligases were recoded, of which three E3 ligases were recoded within defined domains related to ligase activity. Two such recoding events were located in the catalytic ubiquitin-transferring HECT domains for SMURF2 and WWP2, while one recoding event was found in the E2 conjugase RING domain of MARCHF5. MARCHF5 is a mitochondrial E3 ligase involved in the regulation of mitochondrial fusion and fission.^49,50^ We detected an E53G recoding event with high RNA editing frequency that is also conserved across several squid and cuttlefish species (**Fig. 8B**).^15^ Intriguingly this recoding event results in the conversion of the squid sequence to the same amino acid as the homologous position in human MARCHF5 (Gly 57) (**Fig. 8C**), with mutation of this residue to the squid amino acid (G57E) being predicted as pathogenic by AlphaMissense analysis.^51^ Therefore, we hypothesized that this recoding site may influence MARCHF5 activity and/or substrate selectivity.

**Figure 8.**
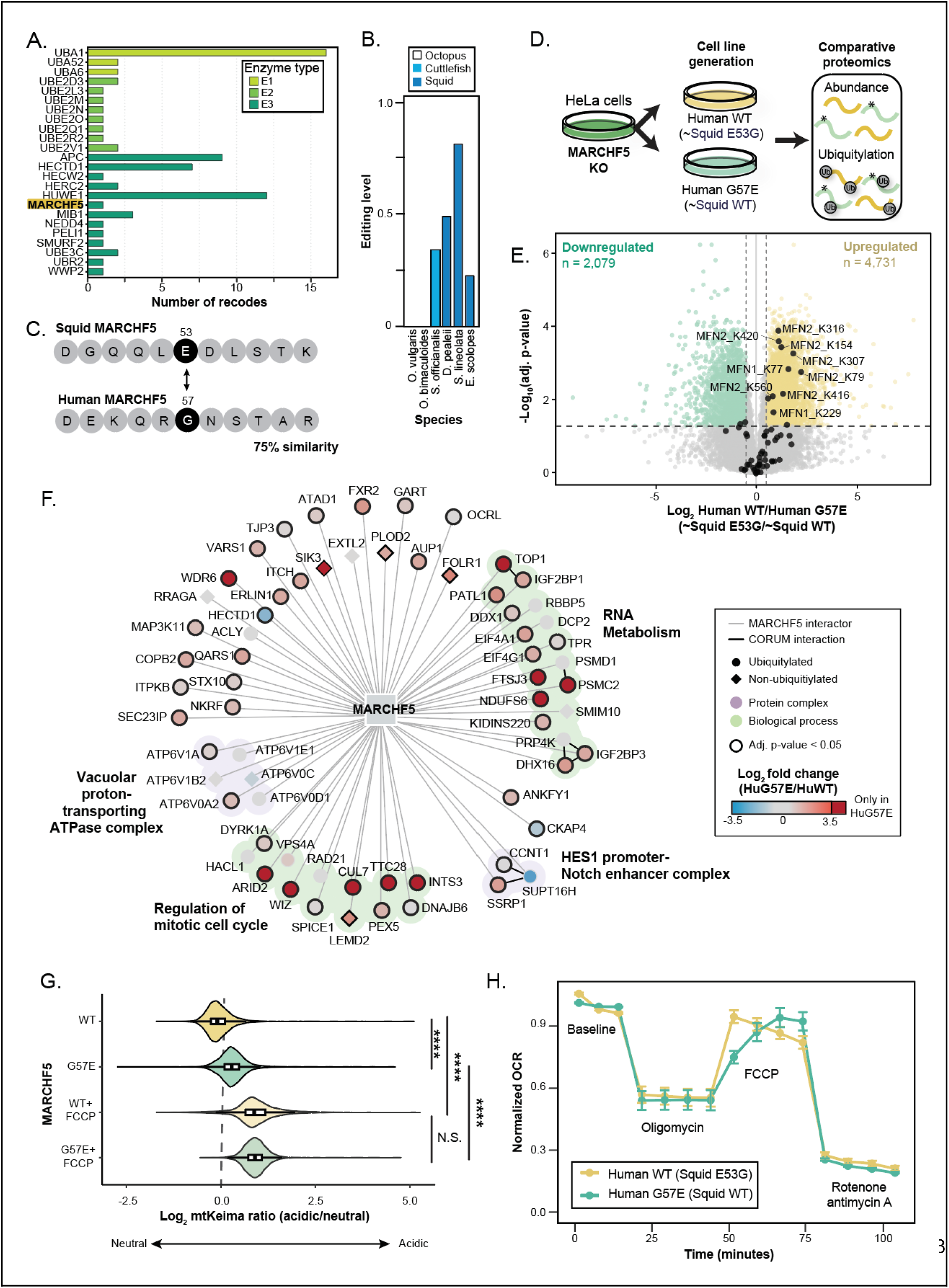
Ubiquitylation signaling by the MARCHF5 E3 ligase is regulated by RNA editing. **A.** Bar graph counting the number of edits identified across ubiquitylation machinery. Proteins are grouped by process (E1-E3), MARCHF5 is highlighted in yellow. **B.** Transcriptomic editing level of MARCHF5 E53G across coleoid species. **C.** Schematic comparison of human and squid MARCHF5 surrounding the E53G recoding site. **D.** Workflow for the generation and enrichment of MARCHF5 complementation cell lines. **E.** Volcano plot comparing ubiquitin site enrichment between the human WT and human G57E mutation. Known substrates of MARCHF5 are marked and labeled in black. **F.** Network diagram of high-confidence (SAINT BFDR < 0.05) MARCHF5 interactions between WT and G57E. Nodes are colored by log2 fold change, node outline denotes significance (adj. P-value < 0.05), node shape denotes if the site was found to be ubiquitylated in panel E. **G.** Violin plot for mt-Keima assay comparing WT and G57E cells with and without FCCP, t-test p-value < 0.05. **H** Seahorse stress test time course assay comparing WT and G57E cells. Dots denote average values (n = 23) with SEM. Multiple time points were taken for each condition. Conditions are noted by text below time point measures.

As there are currently no cephalopod cell lines available, we evaluated this hypothesis in MARCHF5 knock-out HeLa cells using an established complementation assay.^52^ Protein abundance and ubiquitylation sites were measured in cells exogenously expressing either WT human MARCHF5 (mimicking the recoded squid amino acid), or the human MARCHF5 with a G57E mutation (mimicking the WT squid amino acid) (**Fig. 8D**). We found that this single amino acid substitution resulted in dramatic rewiring of ubiquitylation signaling such that 6,450 ubiquitylation sites were significantly regulated (**Fig. 8E, Table S1**). Among these regulated ubiquitylation sites were many known MARCHF5 substrates, such as mitofusin 1 and 2 (MFN1, MFN2), as well as sites diagnostic for various ubiquitin chain structures (**Fig S7A)**. Nearly all ubiquitin chain types were more abundant upon expression of the WT human MARCHF5 protein than the G57E mutation, with the exception of K63-linked chains. To better understand the mechanisms driving such a pronounced phenotype, we performed immunoprecipitation-mass spectrometry (IP-MS) for MARCHF5 in the WT and G57E complementation cell lines (**Fig. 8F**). In total, we identified 66 high-confidence interactors across the two cell lines. Strikingly, these interactions were generally more abundant with MARCHF5 G57E, and almost every interactor was also found to be ubiquitylated in our initial ubiquitin enrichment dataset (**Fig. 8E, Table S1**).

MARCHF5 is an integral membrane E3 ligase residing at the mitochondrial outer membrane that tightly regulates mitochondrial fusion and fission.^49,53^ Thus, we theorized that recoding may alter mitochondrial activity. To test if the single amino acid substitution resulted in functional changes for the cell, we next assayed mitophagy using the pH-sensitive mt-Keima ratiometric reporter^54^ to analyze mitochondrial uptake into acidic lysosomes (**Fig. S7B**). We found that cells expressing either WT or G57E MARCHF5 displayed ramified mitochondria in predominantly neutral pH environments. However, G57E cells (squid WT equivalent) showed a higher degree of mitochondrial acidification, suggesting that G57E increases basal mitophagy (**Fig. 8G**). Importantly, the maximal extent of mitochondrial quality control is comparable across both MARCHF5 variants, as the addition of FCCP, a mitochondrial uncoupler that forces all mitochondria into mitophagy, resulted in a similar mt-Keima ratio for both the WT and G57E forms (**Fig. 8G**). Taken together, these results indicate that cells expressing WT human MARCHF5 have healthier mitochondria and greater relative mitophagic capacity (induced mitophagy compared to basal mitophagy) as compared to those expressing the G57E mutation.

To further investigate the impact on mitochondrial function, we performed a Seahorse assay to measure the cellular oxygen consumption rate of cells expressing either human WT or G57E MARCHF5 (**Fig. 8H**). This measurement is directly proportional to the rate of mitochondrial aerobic respiration, where higher oxygen consumption is associated with a higher electron transport chain activity and indicative of mitochondrial function. To dissect how various mitochondrial contributors affected oxygen consumption, we performed the Mitochondrial Stress Test variation of the Seahorse assay.^55^ In doing so, we found that the Seahorse traces largely yielded comparable results across MARCHF5 genotypes, with the basal respiration rate, ATP production, mitochondrial proton leak, respiratory spare capacity, and non-mitochondrial respiration all being identical between WT and G57E MARCHF5 cells. However, we found that driving maximal respiration via FCCP administration caused a rapid increase in the respiratory rate of WT cells, reaching its maximum rate within 8 minutes. In comparison, this increase was significantly slower in G57E MARCHF5 cells, reaching the maximum respiratory rate after 23 minutes. Because the human WT MARCHF5 is analogous to the E53G recoded squid MARCHF5, these results indicate that recoding of MARCHF5 may enable squid mitochondria to access a healthier mitochondrial state with greater sensitivity and adaptive capacity to mitochondrial uncoupling.

## Discussion

Coleoid cephalopods have long been recognized as an extreme outlier in their use of ADAR-mediated protein recoding. This feature has driven foundational work demonstrating that recoding plays a critical role in protein functions such as neuronal excitability via K^+^ channel gating/assembly, on homeostasis through Na^+^/K^+^-ATPase transport kinetics, and motor protein motility.^21–24^ However, technical challenges have limited our ability to more comprehensively understand the extent to which this recoding is realized at the protein level and its functional consequences. Here, we expand the number of protein-validated recoding sites by more than an order of magnitude and provide a proteome-scale view of how recoding diversifies and regulates protein function in *Doryteuthis pealeii* by altering protein stability, subcellular localization, PTMs, and enzymatic activity. Together, these findings support a model in which cephalopod RNA editing acts not merely as a generator of stochastic proteoform diversity, but as an integral layer of the proteome.

Our global proteomic survey revealed that thousands of RNA-defined recoding events are indeed translated, with 7,998 amino acid substitutions confirmed at the peptide level. The spectrum of recoding types largely mirrors that observed by RNA-seq, with Lys to Arg dominating, and the overall tissue bias in editing (neuronal versus non-neuronal) is recapitulated at the protein level. These benchmarks suggest that the detected sites represent true ADAR activity and are not merely artifacts. At the same time, we detect only ∼36% of the transcriptomically annotated edited sites in our proteomics datasets, and many of those positions appear exclusively as unrecoded WT amino acids. This implies coleoids do not uniformly sample all accessible variants, but rely upon biological filtering between the transcriptome and the proteome, with editing frequency, cell-type specificity, and proteostatic constraints all likely contributing.

Our comparative analyses provide evidence that recoding is selectively tolerated at positions that minimize deleterious structural consequences. First, local alignment to human orthologs combined with AlphaMissense scoring indicates that sites detected only in the recoded form are, on average, predicted to be less pathogenic than sites where only WT or both WT and recoded proteoforms are observed. Second, structural modeling with ESM3 shows that recoded residues tend to result in regions of lower solvent accessibility relative to their WT counterparts. Lastly, sites detected as recoded proteoforms are less likely to be seen across MSAs. These observations suggest that the cephalopod proteome appears to explore an enormous sequence space, but does so under a set of biophysical and evolutionary constraints that filter out the most deleterious options. It appears that the proteome-level realization of RNA editing is biased toward substitutions that are structurally compatible and that avoid strongly destabilizing or aggregation-prone configurations.

Despite this filtering, the realized proteoform space remains extraordinary. Many proteins, including spectrin α-chain and ankyrin-3, exhibit dozens to hundreds of recoding sites at the protein level. For a subset of proteins, certain recoding events are observed only in combination and never individually, implying that co-occurrence of recoding events may be under positive selection or co-selected during translation, folding, or quality control. Nullomer analysis further shows that recoding can generate thousands of short peptide sequences that are absent not only from the unrecoded squid proteome but also from curated proteomes in SwissProt, indicating that RNA editing provides access to previously inaccessible sequence motifs without requiring genomic innovation. This capacity to transiently explore forbidden or disfavored sequence space may be particularly advantageous in organisms that rely on rapid adaptation to changing environments.

Functionally, we demonstrate that recoded proteoforms can differ from their WT counterparts in multiple, orthogonal ways. Solvent-shift profiling reveals that roughly one-third of the analyzed sites measurably change protein stability, with a bias toward destabilization and a broad spectrum of effect sizes. The examples of CenG1A and Loquacious illustrate that even within a single protein, distinct recoding sites—or combinations of recoding sites—can drive stabilizing, neutral, or destabilizing outcomes, implying that functional consequences cannot be inferred from amino acid identity alone. Moreover, our observation that the majority of recoding events do not substantially alter stability suggests that many recoding events modulate other aspects of protein function, such as interaction networks or PTM state.

Our subcellular fractionation experiments show that recoding can rewire protein localization patterns on a proteome-wide scale. The recoding of PBRM1 K577R, located in a putative bipartite nuclear localization signal, provides a striking example in which the recoded proteoform loses nuclear localization and instead accumulates in membrane-associated fractions. This suggests that cephalopods may repurpose proteins, through localization changes, to participate in non-canonical functions through RNA editing. Given that localization is a key determinant of interaction networks, spatial reprogramming provides a powerful way for editing to generate context-dependent phenotypes from a single gene.

A novel insight from this work is the demonstration that ADAR editing can directly regulate the post-translational modification landscape within the proteome. Globally, we identified hundreds of cases in which recoding creates or ablates a PTM site. This manifested most frequently as Lys to Arg recoding, enabling the ablation of acetylation or ubiquitylation. In general, Lys has the most diversified array of PTMs, including SUMOylation, ADP-ribosylation, and numerous types of lipidation. Thus, it may be that Lys to Arg recoded sites ablate other PTMs not covered in our analyses. Conversely, we found that recoding events can create novel phosphorylation sites through Asn to Ser recoding. Although this event was relatively rare, it demonstrates a unique mechanism in which a PTM is dependent on a post-transcriptional modification, providing a novel two-step regulatory system in coleoids. We also observe phosphorylation sites that only appear in the context of a recoding event to be most commonly driven by Ser to Gly recoding proximal to the site of phosphorylation. Similar to what has been observed by DNA mutations in cancer, this can provide an additional mechanism of controlling kinase specificity by which recoding can alter motif elements.^56^ Thus, RNA editing can both directly recode PTM-harboring amino acids, as well as to more subtly remodel the surrounding sequence context that specifies enzyme-substrate interactions. Broadly speaking, this positions ADAR activity as a post-transcriptional regulator of post-translational signaling, allowing it to modulate PTM states without altering DNA or PTM machinery.

Our mechanistic validation of arginine kinase (KARG) and the MARCHF5 E3 ligase illustrates how ADAR-driven recoding manifests within individual proteins. For KARG, we show that multiple recoding events intersect with both phosphorylation and acetylation in a coordinated manner to control enzymatic activity, stability, and protein-protein interactions. The N172S recoding of KARG is one of 19 phospho-creative sites; this event was destabilizing alone, but was further stabilized by phosphorylation, essentially buffering the structural impact conferred from recoding. Interestingly, this recoding site lies within the linker region between the two enzymatic domains, providing a possible explanation for why it does not dramatically impact activity, but does alter protein-protein interactions. Other recoding events within the phosphotransferase domain, e.g., K327R and K352R, can enhance catalytic activity, while acetylation of the unrecorded positions can tune activity up or down, generating a novel regulatory space in which RNA editing and PTMs work in concert to control enzymatic activity. For MARCHF5, we demonstrate that a single recoding event, E53G, within the RING domain is able to reshape ubiquitylation signaling, alter the interactome, and change basal mitophagy. These examples illustrate how coleoids can use RNA editing not only to explore novel protein space, but also to embed recoding events within established regulatory network architectures.

Taken together, our data support a model in which coleoids use ADAR-mediated RNA editing as a multi-dimensional regulatory axis capable of operating across distinct modalities. Rather than acting solely as a mechanism to generate proteoform diversity, editing is selectively realized at positions where its effects can be integrated into a broader cellular context. At the same time, biophysical and functional filters shape which variants can persist and become functionally relevant. The detection of instances in which only the recoded proteoform was identified, despite being evolutionarily rare, raises the possibility that cephalopods have evolved mechanisms to identify and filter against deleterious sites. This model may help to explain how cephalopods can maintain high levels of coding-sequence edits without a complete collapse of proteome integrity.

In the future, using coleoid cephalopods as a biologically encoded means of exploring protein functional space at scale could offer an *in vivo* complement to directed evolution and mutational scanning. Additionally, the ability to control PTM landscapes via RNA editing provides a novel therapeutic strategy for targeting PTMs, such as in the context of tau phosphorylation in neurodegeneration.^57^ This can be further expanded for the targeted control of protein function and localization. Although this study is the most comprehensive proteome investigation to date, it has also raised several new questions. Such as how cephalopods can tune specific edits both inter- and intracellularly, and what quality control mechanisms recognize and manage which edits are faithfully translated.

The current study has limitations that should be addressed. For one, we are only able to capture a piece of the proteoform space that cephalopods occupy with mass spectrometry-based bottom-up proteomics. This limitation will most likely be addressed by future technologies, such as nanopore sequencing of full-length proteins. Additionally, the ability to perturb edits directly in cephalopods is still in its infancy, with the first genetically modified cephalopod only recently being generated.^58^ The lack of cephalopod cell lines, or even closely related model organisms, poses a significant challenge to the mechanistic interrogation of recoding events. Therefore, the development of new models will be essential to understand how the recoding landscape is regulated and deployed.

## Resource availability

All proteomics data have been deposited in the PRIDE repository.^59^

All code used for data analysis has been included as a supplemental file.

## Acknowledgments

We thank Dr. Mariusz Karbowski for providing the MARCH5 KO Hela cell line. We thank Dr. Jesse Rinehart for providing the pSerOTS plasmid and C321 *E. coli* strains. We thank Eli Eisenberg for helpful feedback. We thank the Gladstone Institutes Histology & Light Microscopy Core for supporting the confocal microscopy experiments. We thank Ken Nakamura (Gladstone Institutes) for supporting the Seahorse measurements of mitochondrial respiration. We thank the Cephalopods Program at the Marine Biological Laboratory for providing specimens of *Doryteuthis pealeii*. This work was supported by an NIH grant R01GM133981 (DLS), an NIH Ruth L. Kirschstein National Research Service Award 1F32AI176916-01A1 (JMM), a California Institute of Regenerative Medicine Fellowship (EKK), an NSF grant NSF-BSF:RoL:Imagine 2110074 (JJCR), and a Kavli Foundation Exploration Award: Neurobiology in a Changing Ecosystem (JJCR).

## Author contributions

Conceptualization: JMM, DLS, JJCR. Data curation: JMM. Formal analysis: JMM, MG, EKK, BJP. Funding acquisition: NJK, JMM, DLS. Investigation: JMM, ALR, EKK, RGB, ES. Methodology: JMM, DLS, ALR, PC. Project administration: NJK, DLS. Software: JMM, MG, BJP, DS, MB, BC, PC. Resources: JJCR. Supervision: JJCR, NJK, DLS. Validation: JMM. Visualization: JMM, ALR, EKK. Writing – original draft: JMM, ALR, DLS. Writing – review & editing: all authors.

## Supplemental information

**Table S1.** Contains a list of add recoding sites detected and a variety of properties for them as described across the manuscript, as well as the results of all quantitative proteomics experiments on KARG and MARCHF5.

## STAR Methods

### Specimen collection and dissection

Adult specimens of *Doryteuthis pealeii* were collected by otter trawl in the Vineyard Sound adjacent to Woods Hole, MA. After capture, animals were brought back to the Marine Resources Center of the Marine Biological Laboratory and maintained in large (8ft diameter) circular tanks with flowing natural seawater for up to 48 hrs. Before dissection, squids were euthanized by rapid decapitation. Optic lobes and stellate ganglia were manually dissected in sterile filtered natural seawater, blotted dry, and then flash frozen in liquid nitrogen. For experiments using the proteasome inhibitor MG-132, freshly dissected optic lobes were cut into 4 equal pieces with a razor blade in filtered seawater, and 1/2 of an optic lobe (2 pieces) were used per treatment. Tissue fragments were incubated in squid growth media with 5% FBS in 12-well dishes.^60^ MG-132 was added to the wells at 10 μM or 1 μM, and samples were incubated for 4, 24, or 48 hrs. Control wells had 0.1% or 0.01% DMSO. Following incubations, tissue samples were removed and flash frozen in liquid nitrogen. All protocols using live squid were conducted in accordance with the Marine Biological Laboratory IACUC protocol 22-13A for J Rosenthal.

### Implementation of ADAR-derived recoding sites

ADAR editing sites, frequency of edits, and protein FASTA were taken from previously published results.^17^ The supplement contains the original site data with editing frequency rates in Table S1. Custom R code was used to identify codons that contained multiple edits; an additional editing database was generated from this. The protein FASTA file can be found as Dpealeiiv2.fasta. The amino acid edit tables were used to generate a custom search database.

### Proteoform combinatorial analysis

Proteoform combinatorics were quantified using an R pipeline (data.table, Biostrings, ggplot2) applied to the aggregated multi-protease PSM table. Coefficients of the polynomial were used to obtain the number of proteoforms with exactly 1, 2, or 3 recoded sites (singles, doubles, triples). Observed proteoform counts per protein were defined as the number of unique recoded combinations detected in the PSM data (plus one for the WT form). Proteins were ranked by their theoretical proteoform count; log10 observed versus theoretical counts were plotted to visualize how completely the combinatorial proteoform space was sampled, including split panels highlighting the bottom 99% versus top 1% of proteins by combinatorial capacity. The full script used for this analysis is provided in proteoform_combinatorics.R.

### K-mer Extraction

K-mer extraction and preprocessing: Protein sequence k-mer analyses were performed using the kmerdb_stats software package (https://github.com/Georgakopoulos-Soares-lab/kmerdb_stats, Georgakopoulos-Soares Lab). All analyses used amino-acid–level k-mers composed of the 20 canonical residues (GALMFWKQESPVICYHRNDT). Nullomers were defined as amino-acid k-mers that are possible in principle, but absent from a given proteome. For each FASTA file, nullomers were computed by subtracting the set of observed k-mers from the universal k-mer space. K-mers were then analysed in R to quantify the set differences.

### Orthology inference

Orthology inference was determined via Orthofinder.^61^

### AlphaMissense

Predicted pathogenicity scores for all possible amino acid substitutions were obtained from the AlphaMissense database^32^, which provides a continuous score from 0 to 1 representing the likelihood that a given missense variant is pathogenic. Human UniProt accessions were first normalized to base identifiers (isoform suffixes removed), and only single amino acid substitutions (e.g., A123T) were retained. Squid recoding sites were previously mapped to orthologous human residues using 1:1 OrthoFinder protein pairs and global Biostrings alignments. Only sites where the human reference amino acid matched the WT squid residue were retained. Pathogenicity scores can be found in Table S1.

### ESM3

ESM3 analyses were performed using beta access to the EvolutionaryScale Forge API (model esm3-medium-2024-08) via a custom Python script (ESM3_Structures.py). A private API token was obtained from EvolutionaryScale and used to authenticate all requests. Where PDB coordinates were available, we converted ESM3 outputs to ProteinChain objects (ESM3 SDK + Biotite), and used these to compute (i) per-residue solvent-accessible surface area (SASA), values can be found in Table S1.

### ThermoMPNN

A dockerized version of ThermoMPNN (version: 1:0.0) was used to predict changes in protein stability attributable to recoding events. Protein stability inference was determined from the ‘custom_inference.py’ script with default ThermoMPNN model weights. Input protein PDBs were sourced from ESM3. The ThermoMPNN docker image can be found under martingordon808/thermompnn (https://hub.docker.com/r/martingordon808/thermompnn). ThermoMPNN values can be found in Table S1.

### ConSurf

To assess residue-level amino acid conservation across orthologous proteins, we employed the ConSurf standalone pipeline (v1.0), adapted for high-throughput batch execution. The analysis was run on a custom set of *Doryteuthis pealeii* protein sequences, distributed across multiple batch FASTA files. Each ConSurf run proceeded through the sequential steps. Homology search: MMseqs2 (v13-45111) was used to search for homologs in a local UniRef90 MMseqs2-formatted database. Search parameters included maximum homologs: 100, minimum percent identity: 20, e-value cutoff: 1e-2, coverage threshold: 0.6. MSA and phylogeny: MUSCLE was used to generate a multiple sequence alignment (MSA), and a maximum likelihood phylogenetic tree was inferred from the MSA using ConSurf’s internal pipeline. If fewer than 5 homologs were found, the run was terminated early. Conservation scoring: rate4site (version 2.01) was used to assign a normalized conservation score per residue using the JTT model. The reference sequence (query) was scored based on the evolutionary rate inferred from the tree. Amino acid frequency profiling: For each position in the MSA, the frequency (%) of each standard amino acid (A, C, D, …, Y) was calculated using a custom Python post-processing script. The consurf scores and amino acid frequency information can be found in Table S1.

### Sample preparation for proteomic analysis of recoded peptides

Sample preparation for proteomic analysis of recoded peptides: A detailed breakdown of sample preparation for each dataset is provided in the Supplementary Methods. In general, squid tissues were lysed in trifluoroacetic acid (TFA), followed by 2 cycles of sonication in a Bioruptor system (Hielscher UP200St) for 2 minutes at 100% amplitude per cycle. TFA was quenched by adding 8 volumes of 2 M Tris base (Sigma-Aldrich, T1503). Tris-(2-carboxyethyl)phosphine (TCEP; Thermo Fisher Scientific, 77720) and 2-chloroacetamide (CAA; TCI, C2536) were added to final concentrations of 10 mM and 40 mM, respectively. Samples were incubated in an Eppendorf ThermoMixer C at 45 °C with shaking at 1,100 rpm for 30 minutes. Protein concentration was determined using the Pierce 660 nm Protein Assay (Thermo Fisher Scientific, 22660). Samples were diluted 1:10 in HPLC-grade water and compared against a bovine serum albumin (BSA) standard curve (Thermo Fisher Scientific, 23208). A fixed quantity of total protein was taken forward for each replicate, typically 4 mg per sample.

SP3 bead preparation and protein binding: A Sera-Mag (SP3) bead mixture was prepared by combining Sera-Mag™ Carboxylate-Modified Magnetic Beads (Cytiva 44152105050350 and 65152105050350) at a 1:1 ratio. Beads were washed five times with 2 bead volumes of water and resuspended to the starting concentration. Washed SP3 beads were added to each sample at 100 µL per 0.5 mg protein. Sample volumes were adjusted with water to an equal volume, and 100% ethanol (Koptec Pure Ethanol, 200 proof) was added to a final concentration of 50% (v/v). Samples were incubated in a ThermoMixer at 1,200 rpm for 10 minutes and then placed on a magnetic rack. The supernatant was discarded, and the beads were washed four times with 80% ethanol. For multi-protease digestions, samples were split proportionally according to the number of proteases used and transferred into 1.5 mL Protein LoBind tubes (Eppendorf, 0030108116). Samples were magnetized again, and the supernatant was discarded. Samples for trypsin, LysC, and GluC were resuspended in 400 μL 50 mM triethylammonium bicarbonate buffer (TEAB; Sigma-Aldrich T7408) with protease added at 1:50. Samples for ArgC were resuspended in 400 μL 100 mM HEPES pH 8 (Thermo Fisher Scientific J63578.AP), 10 mM CaCl_2_ (Thermo Fisher Scientific J63122.AE), 50 mM DTT (Sigma-Aldrich D0632), 5 mM EDTA (Thermo Fisher Scientific AM9260G). Samples for ArgC Ultra were resuspended in 20 mM HEPES pH 7.5 (Thermo Fisher Scientific J60712.AP), 10 mM L-Cys (Sigma-Aldrich 1028380100). Samples for AspN were resuspended in 50 mM HEPES pH 8. All digestions were performed overnight (∼16 hours) at 37 °C, shaking at 1,100 rpm.

Post-digestion cleanup and peptide fractionation: Following digestion, samples were transferred to 15 mL conical tubes, and acetonitrile (ACN; Fisher Chemical A955-4) was added to a final concentration of ≥95% ACN to enable SP3 capture for cleanup and fractionation. Samples were incubated with shaking at 1,200 rpm for 10 minutes, placed on a magnetic rack, and the supernatant was discarded. Beads were resuspended in 1 mL of 96% ACN and transferred to 2 mL Protein LoBind tubes. Samples were washed four times with 1 mL 96% ACN. After the final wash, samples were resuspended in 1 mL 96% ACN. From this, 950 µL was transferred to a fresh Protein LoBind tube, magnetized, resuspended in 500 µL 50 mM TEAB, and dried overnight in a speed-vac (LabConco CentriVap). These samples were used for unfractionated peptide analysis. The remaining 50 µL containing peptides was magnetized, and the supernatant was removed. Beads were resuspended in 500 µL of 95% isopropanol (IPA; Fisher BP2618-4), 5% HPLC-grade water, magnetized, and the supernatant collected for fractionation, as previously described.^62^ Sequential elutions were performed using 95%, 90%, 85%, 80%, 75%, 70%, 65%, and 0% IPA, generating eight peptide fractions. Fractions were dried completely in a speed-vac and stored at –80 °C until LC-MS/MS analysis.

### Sample preparation for proteomic analysis of post-translationally modified peptides

Phosphopeptide enrichment: Bulk dried material from the abundance preparations was resuspended in 1 mL 80% ACN, 0.1% TFA. Fe-IMAC beads (Cell Signaling Technology, #20432) were prepared according to manufacturer instructions: 900 µL total bead slurry was washed four times with 1 mL 80% ACN, 0.1% TFA. After washing, the beads were resuspended to the original volume (900 µL), and 25 µL of the washed beads were dispensed into 2 mL Protein LoBind tubes for each enrichment. Samples were centrifuged at 18,213 × g for 5 min, and the clarified supernatant was transferred to tubes containing 25 µL Fe-IMAC beads. Tubes were sealed with Parafilm and incubated on a rotator at 4 °C for 2 hours. After incubation, tubes were placed on a magnetic rack and the supernatant (flow-through) transferred to new 2 mL LoBind tubes. A second round of enrichment was performed by preparing an additional 900 µL of Fe-IMAC beads as above, aliquoting 25 µL into each tube, and adding the collected supernatant. Tubes were sealed and incubated under the same conditions as the first enrichment. While the second enrichment was incubating, beads from the first enrichment were washed three times with 1 mL 80% ACN, 0.1% TFA. Elution buffer was prepared according to manufacturer recommendations: 0.45 mL 28% ammonium hydroxide (Sigma-Aldrich 221228), 2.05 mL water, and 2.5 mL ACN (total 5 mL). Beads from the first enrichment were eluted by adding 50 µL elution buffer, gently mixing every 2–3 minutes for 10 minutes, and magnetizing to recover the eluate. The supernatant was transferred to 1.5 mL Protein LoBind tubes and acidified with 40 µL 20% TFA. Elution was repeated once and pooled with the first eluate. After the second enrichment finished incubating, washing and elution steps were repeated identically. First and second enrichment eluates were kept as separate fractions. All fractions were dried completely in a speed-vac and stored at −80 °C.

Enrichment of acetylated and ubiquitylated peptides: Dried material was enriched using the PTMScan® HS Acetyl-Lysine Motif (Ac-K) or PTMScan® HS Ubiquitin/SUMO Remnant Motif (K-ε-GG) kit (Cell Signaling Technology, #59322). Samples were resuspended in 1.5 mL 1× HS IAP Bind Buffer #1, then centrifuged at 10,000 × g for 5 minutes at 4 °C. The clarified supernatant was transferred into fresh 2 mL LoBind tubes. Antibody-bead slurry was thoroughly resuspended, and 20 µL per replicate was used for enrichment. Beads were washed four times with 1 mL ice-cold PBS, then combined with the soluble sample. Tubes were sealed with Parafilm and rotated at 4 °C for 2 hours. After incubation, samples were briefly centrifuged and placed on a magnetic rack. The supernatant was discarded, and beads were washed four times with 1 mL ice-cold HS IAP buffer, followed by two washes with ice-cold HPLC-grade water. Peptides were eluted using 50 µL 0.15% TFA in a thermomixer at 800 rpm for 10 minutes. Tubes were magnetized, and the supernatant was transferred to 1.5 mL LoBind tubes. Elution was repeated once and pooled. Samples were dried overnight in a speed-vac.

ADP-ribosylation treatment and enrichment: Prior to protease digestion (as described in the abundance workflow), samples were treated with snake venom phosphodiesterase I to cleave ADP-ribosyl modifications ^63^. Sample volumes were adjusted to 1 mL with water, and MgCl₂ was added to 15 mM final concentration. 200 units of snake venom phosphodiesterase I (Thermo Fisher Scientific, 50-203-6470) were dissolved in 400 µL 1 M Tris pH 7.5 (Corning 46-030-CM) supplemented with 15 mM MgCl₂ (Thermo Fisher Scientific AM9530G). The enzyme mixture was divided across samples, delivering approximately 33 units per sample. Samples were incubated in a ThermoMixer at 37 °C, 1,100 rpm, overnight. The following day, samples were prepared the same way as the abundance proteomic samples listed above. Following digestion, the resulting ribose-phosphate remnant was enriched using the same Fe-IMAC phosphorylation enrichment protocol described above.

### Sample preparation for proteomic analysis of affinity/immuno-precipitation peptides

Detailed protocols for each specific precipitation experiment are provided in the Supplementary Methods. A general workflow is outlined below.

Cell and tissue lysis: Lysis buffer (20 mL per batch) was prepared at either 300 mM sodium glutamate (for human cell lysates) or 460 mM sodium glutamate (for squid tissue lysates) (Sigma-Aldrich, AMBH9AD250BF), supplemented with 50 mM HEPES pH 7.5 (Thermo Fisher Scientific, J60712.AP), 1 mM TCEP (Thermo Fisher Scientific, 77720), 5 mM EDTA (Fisher Scientific, MT-46034CI), 0.5% NP-40 (Sigma-Aldrich, 74385), two protease inhibitor tablets (Roche cOmplete Mini, 4693159001). Wash buffer (100 mL) was prepared at 300 mM sodium glutamate for human cells and 460 mM for squid tissue, 50 mM HEPES pH 7.5, and 0.05% NP-40, and chilled on ice. Biological replicates (n = 3) were thawed on ice and resuspended in 1 mL lysis buffer. Samples were sonicated using a probe sonicator (Fisher Scientific Sonic Dismembrator Model 500) at 10% amplitude for 10 seconds, repeated twice. Lysates were cleared by centrifugation (Eppendorf 5430) at 18,213 × g, 4 °C, for 15 minutes. Protein concentrations were quantified using the Pierce 660 nm Protein Assay (Thermo Fisher Scientific, 22660), diluted 1:10 in HPLC-grade water (Fisher Scientific, W64), and compared to a BSA standard curve (Thermo Fisher Scientific, 23208). 2 mg total protein was used for each replicate.

Bead preparation and antibody/bait coupling: For each replicate, 50 µL of Dynabeads™ M-280 Sheep Anti-Rabbit IgG (Invitrogen, 11203D) or Dynabeads™ His-Tag Isolation and Pulldown (Thermo Fisher Scientific, 10103D) was washed four times with 1 mL ice-cold wash buffer using a magnetic rack. Beads were divided into equal fractions. Human cell lysate was incubated with Invitrogen rabbit anti-MARCHF5 (Thermo Fisher Scientific, MA5-57968) or Cell Signaling Technology rabbit anti-MARCHF5 (#19168). Squid lysate was incubated with 100 µg KARG variants (purified protein bait). Bead–antibody/bait coupling was performed on a rotator at 4 °C for 1 hour. After coupling, beads were placed on a magnetic rack, washed three times with 1 mL ice-cold wash buffer, and resuspended in 600 µL wash buffer.

Immunoprecipitation: 50 µL of prepared beads was added to each clarified lysate (2 mg protein). Tubes were sealed with Parafilm and incubated overnight at 4 °C on a rotator. The following day, the beads were magnetized and the supernatant removed. Beads were washed three times with 1 mL ice-cold wash buffer. Proteins were eluted by adding 100 µL of 0.1 M sodium citrate (Sigma-Aldrich S4641), pH 2.5, and incubating at 800 rpm for 10 minutes in an Eppendorf ThermoMixer C. Elution was repeated one additional time, and eluates were combined. Eluted samples were neutralized by adding 200 µL of 0.5 M HEPES pH 8. Neutralized eluates were processed for digestion using the SP3 sample preparation workflow described for abundance proteomics.

### Solvent shift proteomics

Sample preparation: Gill and optic lobe samples (n = 3 biological replicates each) were pulverized using a cryogenic mill (SPEX SamplePrep Freezer/Mill) at 10 cycles per second, 2 minutes on / 2 minutes off, for 10 cycles. Pulverized material was resuspended in 500 µL lysis buffer composed of 20 mM Tris-HCl (Corning 46-030-CM), 150 mM NaCl (Thermo Fisher Scientific AM9760G), 300 µL 5 M NaCl, 200 µL 1 M Tris, 1 cOmplete Mini Protease Inhibitor tablet (Roche 4693159001), 1 PhosSTOP tablet (Roche 4906845001) and water to 10 mL. All buffers were kept ice-cold. Suspensions were rotated for 30 minutes at 4 °C, then clarified by centrifugation at 750 × g for 2 minutes at 4 °C. Protein concentration was estimated using the Pierce Bradford Protein Assay (Thermo Fisher Scientific 23200) against a BSA standard curve (Thermo Fisher Scientific 23208).

Solvent-shift treatment: Each lysate was divided into ten 40 µL aliquots. A solvent mixture (AEA) consisting of 50% acetone (Koptec V1825), 50% ethanol (Koptec 200-proof ethanol), and 0.1% acetic acid (Sigma-Aldrich 320099) was added to generate final concentrations of 0–27% AEA (0, 3, 6, 9, 12, 15, 18, 21, 24, 27%). AEA was added along with lysis buffer to a final volume of 80 µL, followed by an additional 20 µL of lysis buffer, yielding 100 µL total volume per condition. Samples were vortexed briefly and centrifuged at 21,000 × g for 45 minutes at 4 °C. 40 µL of each soluble fraction was transferred into a 96-well plate (Thermo Fisher Scientific AB-0800 277143).

Denaturation, reduction, and alkylation: A 10 mL denaturation stock was prepared by dissolving 1.2 g urea (Sigma-Aldrich U5378) and 1.2 mL 1 M Tris, bringing the volume to 10 mL with water. 40 µL of this denaturation buffer was added to each well, followed by 8 µL of a 100 mM TCEP / 400 mM CAA mixture (Thermo Fisher Scientific 77720; TCI C2536). Samples were incubated at 37 °C, 800 rpm for 5 minutes using an Eppendorf ThermoMixer C. Because solvent treatment lowered pH, each reaction was adjusted to pH 7–8 using 0.1 N NaOH (Sigma-Aldrich 2105). Samples were then split equally into two identical 96-well plates (44 µL per well). One plate was used for trypsin/LysC digestion and the other for GluC digestion, with each well receiving 2 µg sequencing-grade trypsin (Promega V5113) and 1 µg LysC (Wako 125-02543, 2 µg/µL) or 2 µg GluC (Promega V1651, 2 µg/µL). Samples were digested overnight at 37 °C, 500 rpm, followed by an additional 1 µL trypsin or 1 µL GluC for 1 hour at 37 °C the following morning.

### LOPIT-DC proteomics

Sample preparation: Gill and optic lobe tissues (n = 3 biological replicates per tissue) were thawed on ice in the cold room and lysed in LOPIT buffer consisting of 0.25 M sucrose (Sigma-Aldrich S1888), 10 mM HEPES pH 7.5 (Thermo Fisher Scientific J60712.AP), 460 mM mono-sodium glutamate (AmBeed - A749276), 2 mM EDTA (Thermo Fisher Scientific AM9260G), 2 mM magnesium acetate (Sigma-Aldrich M2545), 1 cOmplete Mini protease inhibitor tablet (Roche 4693159001) Lysis buffer was prepared in 10 mL total volume using: 100 µL 1 M HEPES, 4.27 mL 20% sucrose, 40 µL 0.5 M EDTA, 40 µL 0.5 M magnesium acetate, and water to volume. Approximately 2 mg total protein was obtained per replicate. Lysates were kept on ice at all times. To remove unlysed debris, samples were centrifuged at 200 × g for 5 minutes at 4 °C. Pellets from this clarifying spin were retained and stored at −80 °C.

Differential centrifugation workflow: Subcellular fractionation was performed using the LOPIT-DC workflow^42^ at 4 °C, using pre-cooled tubes and keeping all fractions on ice. After each centrifugation, the supernatant was transferred to a fresh pre-chilled tube for the next spin, while pellets were kept on ice. Sequential centrifugation steps (g-force × time) were as follows:

1. 1,000 × g, 10 min - Pellet 1
2. 3,000 × g, 10 min - Pellet 2
3. 5,000 × g, 10 min - Pellet 3
4. 9,000 × g, 15 min - Pellet 4
5. 12,000 × g, 15 min - Pellet 5
6. 15,000 × g, 15 min - Pellet 6
7. 30,000 × g, 20 min - Pellet 7
8. 79,000 × g, 43 min - Pellet 8
9. 120,000 × g, 45 min - Pellet 9

The final post-ultracentrifugation supernatant was precipitated by adding ice-cold methanol to 90% final concentration, vortexing briefly, and centrifuging at 5,000 × g for 5 minutes at 4 °C to generate Pellet 10. Pellets were visibly larger in gill samples compared to the optic lobe, consistent with increased particulate content.

Pellet solubilization, reduction/alkylation, and digestion: Each pellet was resuspended in 20 µL 8 M urea / 100 mM ammonium bicarbonate (ABC) pH 8 (prepared as: 4.8 g urea + 1 mL 1 M ABC, water to 10 mL). Reduction and alkylation were performed by adding 1:10 volume of freshly thawed TCEP/CAA stock consisting of 100 mM TCEP (Thermo Fisher Scientific 77720) and 440 mM chloroacetamide (CAA) (TCI C2536). The stock was prepared by dissolving 143.3 mg TCEP + 187.0 mg CAA in 5 mL HPLC-grade water. 2 µL of TCEP/CAA mix was added to each 20-µL pellet suspension, followed by incubation at 37 °C for 45 minutes at 500 rpm in a ThermoMixer C (Eppendorf). For enzymatic digestion, each sample was diluted 8-fold with 100 mM ABC pH 8. An enzyme master mix was prepared by combining 1.2 mL 100 mM ABC, six vials of sequencing-grade trypsin (Promega V5113; 0.5 µg/µL), and 20 µL LysC (Wako 125-02543; 2 µg/µL). 20 µL of enzyme mix was added to each fraction (10 fractions per replicate), and digestion proceeded overnight at 37 °C, 300 rpm. Peptides were desalted using 96-well C18 plates (The Nest Group Inc., HNS S18V; 20 mg PROTO 300 C18 resin per well). Samples were acidified to pH 3 with 10% TFA (Sigma-Aldrich 302031), clarified by centrifugation at maximum speed for 5 minutes, and loaded onto C18 plates pre-wetted with 400 µL 80% ACN + 0.1% TFA. After loading, plates were washed three additional times with 0.1% TFA. Peptides were eluted twice with 55 µL 50% ACN + 0.25% formic acid (FA) (Thermo Fisher Scientific LS118). Combined eluates were dried by speedvac (LabConco CentriVap).

### Mass spectrometry data acquisition

All samples were resuspended in 2% ACN, 0.1% FA (Thermo Fisher Scientific LS118) with a 50 µL volume for abundance samples and 25 µL volume for PTM-enriched samples. Peptide concentrations were estimated using A205 measurements on a NanoDrop One spectrophotometer (Thermo Fisher Scientific) using the standard A205 with the 31-correction method. Where possible, samples were adjusted to 250 ng/µL. A 0.45 µm filter plate (Sigma-Aldrich MSHVN4550) was pre-conditioned with 25 µL 2% ACN, 0.1% FA, and centrifuged at 500 × g for 10 minutes. Samples were then loaded and filtered under the same conditions.

LC-MS/MS data acquisition: Unless otherwise specified, acquisition parameters were similar across all sample types. Detailed instrument files for each dataset are provided in the supplement and PRIDE upload. 500 ng of digested peptide was injected and analyzed with an Orbitrap Exploris 480 mass spectrometry system (Thermo Fisher Scientific), equipped with a Vanquish Neo UHPLC and a Nanospray Flex ion source. Chromatographic separation was performed using a PepSep C18 column (Bruker 1893474; 15 cm × 150 µm, 1.5 µm particle size) fitted with a PepSep stainless steel emitter (Bruker 1893525). Mobile phases were A: 0.1% FA in water and B: 0.1% FA in 80% ACN (Thermo Fisher Scientific LS122). In general, peptides were separated with the following gradient: 2% to 5% B over 2 min, 5% to 25% B over 80 min, 25% to 40% B over 10 min, 40% to 95% B over 2 min, and held at 95% B for 6 min. The column was then returned to initial conditions and equilibrated prior to the next injection.

Mass spectrometer settings: Data-dependent acquisition (DDA) was used for the majority of runs, with the exception of the solvent shift and LOPIT data. Full settings for each experiment can be found in the supplement and are uploaded with the PRIDE data. The DDA parameters are provided in full; MS1 (Full Scan), m/z range: 350–1150, resolution: 120,000 (at m/z 200), AGC target: 300% (normalized), RF lens: 50%, max injection time: Auto, scan mode: Orbitrap. dynamic exclusion: 20 s, exclusion mass tolerance: ±10 ppm. MS2 (HCD Fragmentation): Precursor charge selection: 2–6, isolation window: 1.6 m/z, HCD collision energy: 28%, AGC target: 300%, resolution: 30,000 (at m/z 200), max injection time: 54 ms.

### Mass spectrometry data analysis

Tesorai database search: Tesorai^64^ employs a deep learning scoring algorithm trained on over 100 million real peptide–spectrum matches from multiple search engines and uses a novel decoy-generation strategy that improves false discovery rate (FDR)^65^ control compared to on-the-fly approaches such as MSFragger.^66^ Given its robustness and high-fidelity scoring, Tesorai was selected as the primary search engine for all multi-protease abundance datasets and all phosphorylation and acetylation enrichment proteomics. Search settings included: minimum peptide length: 7 amino acids, maximum missed cleavages: 2, variable modifications: Oxidation (M), N-terminal acetylation, Deamidation (N/Q), Phosphorylation (S/T/Y) as needed, Lys acetylation (K) as needed; fixed modification: Carbamidomethyl (C). The FDR was set to 1% at PSM-level post-processing. Each protease dataset was searched against a custom FASTA file.

FragPipe search for ubiquitin-modified peptides: Because Tesorai did not support the identification of ubiquitylated peptides at the time of analysis, ubiquitin remnant (K-ε-GG) datasets were analyzed using FragPipe v22.0 with MSFragger and Philosopher.^67^ Full parameter files (fragger.params, workflow XMLs) are provided on PRIDE. Settings (modifications and digestion rules) included: Enzyme: Trypsin (K/R, not before P), missed cleavages: 2, variable modifications: Oxidation (M, +15.9949; max 3 per peptide), N-terminal acetylation (+42.0106; max 1 per peptide), Deamidation (N/Q, +0.984016; max 3 per peptide), Ubiquitin remnant (K), +114.04293; max 1 per peptide and fixed modification: Carbamidomethyl (C, +57.02146). Maximum variable modifications per peptide: 3 with a peptide length of 7 to 50 amino acids. A custom FASTA was generated as above with decoys and contaminants appended by FragPipe. Default FragPipe workflow settings were used unless otherwise noted.

DIA/Spectronaut analysis for solvent-shift and LOPIT: Solvent shift and LOPIT data independent acquisition (DIA) data were analyzed using Spectronaut^68^ 19.8.250311.62635. Spectral library generation: A DDA-based spectral library was first generated from representative DDA runs. The library (provided as TSV files in PRIDE) was constructed using: enzyme: Trypsin/P, variable modifications: Oxidation (M), N-terminal acetylation, deamidation (N/Q), and fixed modification: Carbamidomethyl (C). FDR: 1% at PSM, peptide, and protein levels (BGS Factory Settings) with background imputation. DIA analysis: DIA runs were analyzed using the above spectral library and the same FASTA file. Default Spectronaut settings were used except for background signal imputation, which was adjusted as described in ExperimentSetupOverview.txt. Outputs included Results.tsv and MSstats.tsv, deposited in PRIDE.

Downstream statistical analysis in R: All custom analyses were performed in R 4.5.0, and full scripts are provided in the Supplement. PTM and recoded site matching: For abundance, phosphorylation, acetylation, and ubiquitin datasets, peptide and site-level matches to RNA edits or PTMs were extracted using data.table, dplyr, tidyr, and Biostrings. MSstats^69^ was used to analyze search outputs for AP-MS data; general analysis code can be found on the Krogan lab GitHub (https://github.com/kroganlab) and is also provided as usage.spectronaut2ArtMS.R with helper functions in spectronautFile2ArtMS.R. The results-mss-ProteinLevelData.txt file was further processed into inter, bait, and prey files for analysis with SAINTexpress^70^ for MARCHF5 IP-MS data. For KARG AP-MS, differential protein abundance between each mutant bait and WT was assessed on the log_2_ scale using limma’s linear models with contrasts (variant − WT) and empirical Bayes variance moderation, yielding moderated t-statistics and associated p values with a BH correction. To account for variation across replicates, we included a CV cutoff of 30%.

### Solvent shift analysis

MSstats Protein-Level Input and Pre-Processing: Raw DIA data were analyzed using Spectronaut (Biognosys) and processed through the MSstats workflow (usage.spectronaut2ArtMS.R in artMS) to generate protein-level abundance tables (results-mss-ProteinLevelData.txt). Only proteins with a single identifier were retained. Analyses were restricted to samples belonging to the Optic Lobe solvent-shift experiments, each of which contained three biological replicates. Protein-level abundance measurements derived from solvent-shift experiments were normalized and filtered prior to analysis. Stability-related features were extracted from abundance profiles across solvent conditions and used to generate comparative stability metrics. Differences between wild-type and recoded protein forms were evaluated using statistical models.

Comparison of ThermoMPNN ΔΔG predictions with solvent-shift ΔTm50 measurements: ThermoMPNN stability predictions were compared directly to experimentally measured solvent-shift ΔTm₅₀ values.:

### LOPIT analysis

LOPIT analysis: Raw fractionation intensities were obtained from the results-mss-groupQuant.txt file generated by MSstats.^71^ Entries containing multiple protein assignments (Protein IDs containing “;”) were removed. Only Optic Lobe (OL) fractions were used for this analysis. Proteins were classified as WT or recoded based on the presence of the substring _edit_ in the Protein identifier. For recoded entries, the underlying protein base, the recoding string (e.g., N139D_R141G), and the corresponding amino-acid window (e.g., pos126-142) were parsed directly from the FASTA-style identifier. Start and end coordinates were extracted when applicable. LOPIT fraction intensities were supplied as log_2_ values. Each value was converted back to linear space, summed per protein, and normalized to percent of total signal per protein. This produced a 10-fraction distribution (OL_01-OL_10) in which each row sums to 100%. These normalized percent distributions were used for all downstream comparisons and visualization.

Gene and orthology annotation: To annotate each squid protein with a gene-level name, the Dpealeii_Genome_Conversion_Table.csv file was joined to the matrix using the parsed protein_base field. When available, Gene_Name was assigned as the final annotation; otherwise, the genome assembly name was used.

Data restructuring and comparison: Normalized LOPIT values were reshaped to long format to generate a table containing one row per (Protein x Fraction). From this dataset, two filtered tables were generated: (1) WT-only entries for gene-set enrichment analysis, and (2) paired WT-recoded datasets containing only protein families for which both a WT form and at least one recoded variant were observed. For each recoded proteoform, WT and recoded profiles were combined and visualized as line plots showing percent distribution across OL_01-OL_10. Plots were saved both as an aggregated multi-page PDF and as individual SVG files for figure assembly.

GSEA: LOPIT percent-intensity data (WT proteins only) were imported from the row-normalized matrix described above (dt.long.csv). For enrichment, we used protein–annotation terms derived from STRING v12.0; specifically, we restricted those entries annotated as “Component” and built term–to–gene mappings by extracting STRING protein IDs and stripping the species prefix. To identify groups of proteins with similar fractionation behavior, we converted the long table to a matrix and set missing values to zero. We then applied k-means clustering (k = 10) to the percent profiles, using Euclidean distance on the 10-fraction vectors. Cluster membership (cluster ID plus gene identifier) was passed to custom R-scripts available at https://github.com/kroganlab (bp_utils, fastEnrich in enrichmentTestFunctions.R), which perform hypergeometric over-representation analysis of STRING “Component” terms within each cluster relative to the set of all clustered proteins. Resulting p-values were adjusted for multiple hypothesis testing (Benjamini–Hochberg), and enriched terms were ranked by adjusted p-value within each cluster.

LOPIT per fraction statistical analysis: Replicate-level protein intensities were exported from MSstats as the sampleQuant table (results-mss-sampleQuant.txt). Columns corresponding to the optic lobe (OL) fractions (OL_01-OL_10) were retained and reshaped to long format with explicit Fraction (OL_01-OL_10) and Rep (replicate 1-3) identifiers. Proteins were classified as WT or recoded based on the presence of the _edit_ tag in the protein identifier. MSstats log_2_ intensities were back-transformed to linear space and, for each protein and replicate, intensities were normalized to sum to 100% across OL fractions to yield a per-replicate fractional abundance profile (percent). Only proteins for which both a WT and at least one recoded variant were detected were retained. For each recoded variant and each fraction, we compared the replicate-level normalized abundances between recoded and WT using Welch’s two-sample t-test (unequal variances). Tests were only performed when both groups had at least two finite replicate values and non-zero combined variance; otherwise, no p-value was assigned. For each fraction, we recorded the mean WT and recoded percentages, their difference (Δ% = mean_recoded - mean_WT), the t statistic, the nominal p-value, and Cohen’s d effect size. Multiple testing correction was applied within each fraction across all variants using the Benjamini–Hochberg procedure to obtain fraction-wise FDR values (adj_p_by_fraction).

Integration with STRING nuclear protein–protein interactions: High-confidence protein–protein interactions were obtained from STRING v12.0 (STRG0A89UHV.protein.links.v12.0.txt). STRING protein IDs were mapped to protein identifiers by stripping the database prefix, and interactions were treated as undirected edges; when multiple edges existed for the same pair, the maximum combined score was retained. Only interactions with a combined score ≥ 600 were used for downstream analysis. Subcellular localization annotations were taken from the STRING enrichment terms file (110092772.protein.enrichment.terms.v12.0.txt). Proteins were labeled as “nuclear” if any associated Gene Ontology term or annotated keyword contained nucleus-related descriptors. Interactors were restricted to those proteins detected in the LOPIT dataset, and, for each seed protein, we counted both the total number of detected interactors and the number of interactors annotated as nuclear. For the t-test results, we extracted fraction 9 (OL_09) entries from values calculated above, keeping one row per recoded variant with its fraction-specific Δ% (Recoded - WT) and FDR (adj_p_by_fraction). These data were merged with the nuclear PPI counts.

### Plasmid and strain construction

Plasmid construction: A Sleeping Beauty transposon system^72^ was used to integrate doxycycline-inducible MARCHF5 into mammalian cells. The pSBtet-Blast plasmid (Addgene #60510) was grown in 5 mL LB + 100 µg/mL ampicillin (Thermo Fisher Scientific J63807.09) overnight at 37 °C and isolated using a Qiagen QIAprep Spin Miniprep Kit (27106). The plasmid was digested with SfiI and purified using a Qiagen QIAquick PCR Purification Kit (28194). MARCHF5 was PCR-amplified from Addgene plasmid #62039 using KAPA HiFi HotStart ReadyMix (2x) (Roche KK2602). Primers contained 20–30 bp overlaps for Gibson assembly into pSBtet-Blast. Mutagenesis primers were included during PCR to directly assemble the E53G variant. Complete plasmid assembly schematics, sequencing validations, and primers are provided in the supplemental SnapGene files.

KARG plasmids were generated using an E. coli codon-optimized gBlock (IDT) containing 20–30 bp overlaps for Gibson assembly. The protein expression vector (Mohler et al., 2023) was digested with KpnI and SacI, and the KARG insert was assembled by Gibson cloning. The resulting WT vector was used as a template for site-directed mutagenesis to generate N172S, N172TAG, K327R, K327TAG, K352R, K352TAG. Here, TAG indicates replacement of the native codon with the UAG amber stop codon for genetic code expansion. Additional construction history is available in the SnapGene files.

Strain construction: KARG strains were generated in the following backgrounds: C321.ΔA.exp,^73^ C321^XPS^,^47^ and BL21 (New England Biolabs C2530H). Strains were grown overnight in 5 mL LB (Fisher Scientific BP1427-500) at 37 °C, shaking at 230 rpm, without antibiotics. The following morning, cultures were diluted 1:100 into 20 mL LB (one tube each for C321^XPS^ and C321.ΔA.exp; two tubes for BL21) and grown at 37 °C to OD₆₀₀ ∼ 0.8. Cells were chilled, centrifuged at 6,780 × g for 2 min at 4 °C, and washed three times with ice-cold water. After the final wash, cells were resuspended in 1 mL ice-cold water, transferred to a 2 mL microcentrifuge tube, pelleted again at 6,780 × g, and finally resuspended in 100 µL ice-cold water. 50 µL of competent cells were aliquoted per transformation. 100 ng total plasmid (50 ng of each plasmid when 2 are used). DNA was added according to the following combinations:

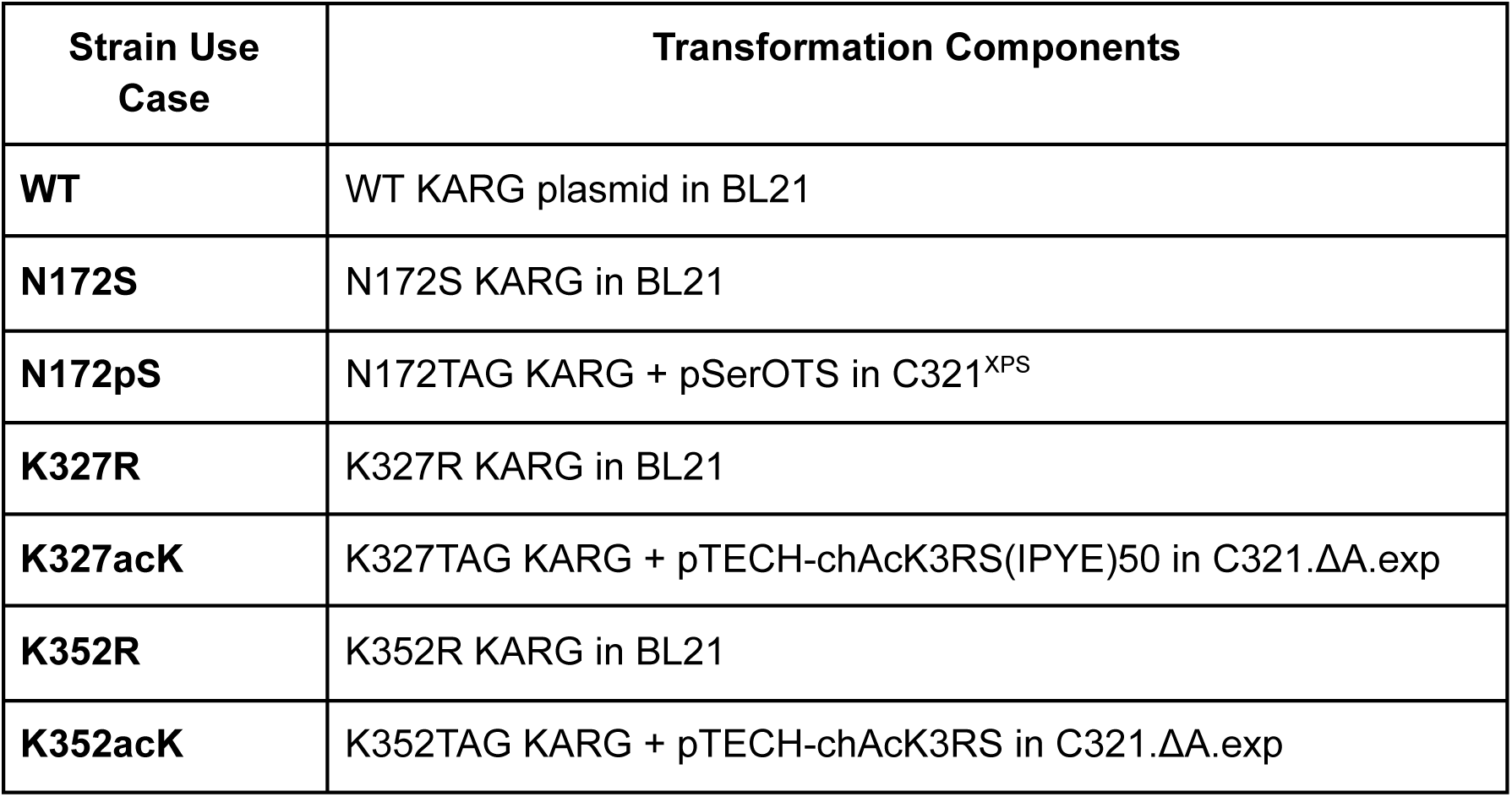

The pSerOTS, C321.ΔA.exp, and C321^XPS^ strains were provided by Dr. Jesse Rinehart, and pTECH-chAcK3RS(IPYE) was obtained from Addgene (#104069). Cells and DNA were mixed gently and transferred to 0.1 cm Gene Pulser cuvettes (Bio-Rad #1652089). Electroporation was performed with a Bio-Rad GenePulser Xcell using preset conditions for E. coli (1 mm cuvette, 1.8 kV, 25 µF, 200 Ω). Immediately after electroporation, cells were resuspended in 600 µL LB, transferred to 1.5 mL tubes, and incubated at 37 °C for 1 hour with shaking. Cells were plated on LB-agar with appropriate antibiotics: 100 µg/mL ampicillin (KARG expression vector), 50 µg/mL kanamycin (Sigma-Aldrich 60615-5G) for pSerOTS, 25 µg/mL chloramphenicol (Sigma-Aldrich C0378) for pTECH-chAcK3RS. After overnight incubation, 10 colonies per construct were picked and grown overnight at 37 °C with shaking. Glycerol stocks were prepared the following morning and stored at −80 °C.

### Expression and purification of arginine kinase

Protein expression: Starter cultures (50 mL LB + appropriate antibiotics) were prepared from glycerol stocks of the KARG strains described above and grown overnight at 37 °C, 230 rpm. The next morning, 9 L LB + antibiotics was inoculated to an initial OD₆₀₀ ∼ 0.01 for each strain. For acetyllysine-containing strains (K327acK and K352acK), cultures were supplemented at inoculation with 3 mM Nε-acetyl-L-lysine (Thermo Scientific J64129.06). Cultures were grown at 37 °C to OD₆₀₀ ∼ 0.8 and induced using 20% arabinose (Sigma-Aldrich A3256-100G) to a final concentration of 0.2%. Acetyllysine-containing strains were additionally supplemented with 20 mM nicotinamide (Sigma-Aldrich N3376) (from a 2 M stock). Cultures were shifted to 16 °C, shaking at 200 rpm, and expressed overnight (∼20 hours). Cells were harvested by centrifugation at 6,000 × g, 4 °C, 10 minutes using 1.5 L flasks (Avanti J-26S XP Centrifuge; JLA 9.1000 rotor). Pellets were resuspended in 120 mL ice-cold LB, split into four 50 mL Falcon tubes, and centrifuged again at 6,780 × g, 4 °C, 20 minutes. Supernatant was removed, and pellets were stored at −80 °C.

Cell lysis: A total of 70 mL lysis buffer per sample was prepared containing: 50 mM Na₂HPO₄ pH 8 (Sigma S7907 dibasic / Acros Sodium Phosphate, Monobasic Monohydrate), 300 mM NaCl (Corning 46-032-CV), 10 mM imidazole (Fisher Scientific O3196-500). 1 mM MgCl₂ (Invitrogen AM9530G), 1 mg/mL lysozyme (Thermo Fisher Scientific 89833). 1 tablet PhosSTOP per 10 mL (Sigma-Aldrich 4906845001), 1 mM PMSF (Thermo Fisher Scientific 36978), 1 mM TCEP, 1 tablet cOmplete Protease Inhibitor per 10 mL (Sigma-Aldrich 4693116001), 125 U/mL benzonase (Sigma-Aldrich E1014-5KU). Phosphate buffer (pH 8.0) was generated by combining 1 M Na₂HPO₄ and 1 M NaH₂PO₄ at the appropriate dibasic:monobasic ratio to yield a final concentration of 50 mM total phosphate. Additional supplements included: 1 mM Na₃VO₄ (Thermo Fisher Scientific J60191.AD) and 20 mM NaF (Sigma-Aldrich 201154) for N172pS samples; 20 mM nicotinamide for K327acK and K352acK samples. Frozen pellets were thawed and resuspended in 60 mL lysis buffer in a 100 mL container. Suspensions were incubated at 37 °C for 10 minutes, vortexed briefly, and sonicated using a probe sonicator at 40% amplitude, 10 sec on / 20 sec off, for a total sonication time of 60 seconds. Lysates were transferred to 30 mL centrifuge tubes (Thermo Scientific 3119-0050PK) and centrifuged at 40,000 × g, 4 °C, 25 minutes. Supernatants were transferred to clean tubes and centrifuged again under identical conditions. Clarified lysates were pooled into 50 mL Falcon tubes.

Purification buffers: Ni-NTA wash buffer (200 mL/sample): 50 mM Na₂HPO₄, 300 mM NaCl, 50 mM NaF, 1 mM TCEP, 20 mM imidazole, pH 8. Ni-NTA elution buffer (15 mL/sample): 50 mM Na₂HPO₄, 300 mM NaCl, 250 mM imidazole, pH 8. 2x Amylose dilution buffer (20 mL/sample): 40 mM HEPES pH 7.5, 2 mM EDTA. Amylose wash buffer (200 mL/sample): 20 mM HEPES pH 7.5, 1 mM EDTA, 200 mM NaCl. Amylose elution buffer (15 mL/sample): 20 mM HEPES pH 7.5, 1 mM EDTA, 200 mM NaCl, 10 mM maltose (Sigma M5885). 2x storage buffer (15 mL/sample): 50 mM HEPES pH 8, 100 mM KCl (Thermo Fisher P217), 20 mM MgCl₂, 20 mM β-mercaptoethanol (Sigma-Aldrich M3148).

Protein purification: During lysate clarification, a gravity-flow Ni-NTA column was prepared with 3-4 mL Ni-NTA resin (50% slurry) (Qiagen Ni-NTA agarose) and equilibrated with 20 mL wash buffer. Clarified lysate was applied to the Ni-NTA column twice. The column was washed with 150 mL Ni-NTA wash buffer, and the bound protein was eluted with 10 mL Ni-NTA elution buffer. The 10 mL Ni-NTA eluate was mixed with 10 mL 2x amylose dilution buffer. An amylose column was prepared with 6 mL amylose resin (3 mL bed volume) (NEB E8021L) and equilibrated with 30 mL amylose wash buffer. The diluted Ni-NTA eluate was applied to the amylose column twice, followed by a 150 mL amylose wash. Protein was eluted with 10–15 mL amylose elution buffer. Eluates were concentrated to 500 µL in Amicon Ultra-4 30 kDa filters (Sigma-Aldrich UFC803008). Three rounds of buffer exchange into 2x storage buffer were performed at 6,000 × g, 4 °C, 10 minutes per spin (increased as needed to reach 500 µL final volume). Purity and size were evaluated by SDS-PAGE and Coomassie staining. Protein concentration was measured by A280 on a NanoDrop using storage buffer as a blank. Finally, an equal volume of 80% glycerol (Fisher Scientific MP113055034) was added to achieve a 40% glycerol final concentration, and aliquots were stored at −80 °C.

### Arginine kinase activity assays

Percent activity assay (ActiveX ATP desthiobiotin probe): To ensure equal amounts of active KARG enzyme were used in downstream ADP-Glo assays, the ActiveX ATP desthiobiotin probe (Thermo Fisher Scientific 88311) was used to quantify the accessible ATP-binding active site for each purified variant. The goal of this assay is to determine the fraction of catalytically competent enzyme by measuring probe binding saturation. The assay was performed using a probe titration series in large molar excess over the enzyme to ensure plateau detection.

ADP-Glo activity assay: A reaction master mix was prepared fresh for each KARG variant containing: 12.5 nM active kinase, 12.5 mM L-arginine, 25 mM HEPES pH 8, 50 mM KCl, 10 mM MgCl₂, 10 mM β-mercaptoethanol. 20 µL of reaction mix was aliquoted in triplicate into a 96-well half-area plate for seven time points: 5, 15, 30, 60, 90, 120, and 180 minutes. Plates were equilibrated at 20 °C for 5 minutes in a ThermoMixer.

A reverse time course was performed by adding 5 µL of 500 µM ATP in 25 mM HEPES pH 8, 50 mM KCl, 10 mM MgCl₂, 10 mM β-mercaptoethanol at each time point; Row H (180 min): ATP added at t = 0, Row G (120 min): ATP added 1 hour later, Row F (90 min): ATP added 30 minutes after that, and continuing in this fashion until the 5-minute time point. After the final (5-minute) time point had incubated for 5 minutes, 25 µL ADP-Glo reagent was added to quench remaining ATP. Samples were incubated at 20 °C for 40 minutes. Next, 50 µL kinase detection reagent was added, followed by a 30-minute incubation at room temperature. Luminescence was recorded using a SpectraMax iD5 plate reader.

### Generation of MARCHF5 cell lines

MARCHF5 -/- HeLa knockout cells were maintained in Gibco DMEM (Thermo Fisher Scientific 11995065) supplemented with 10% FBS (Thermo Fisher Scientific A5209501) and 1% Penicillin–Streptomycin (5,000 U/mL) (Thermo Fisher Scientific 15070063). Cells were seeded in 10 cm dishes and grown to ∼60% confluence. For transfection, the media was replaced with 6 mL serum-free DMEM, and cells were returned to the incubator (37 °C, 5% CO₂) for 1 hour. During this incubation, the Sleeping Beauty transposon and transposase DNA mixture was prepared. A DNA mixture consisting of: 15 ng transposase plasmid (Addgene #34879), 30 ng WT or G57E MARCHF5 pSBtet-Blast plasmid, 5 µg salmon sperm DNA (Thermo Fisher Scientific 15632011) was combined with 250 µL DMEM. Separately, 15 µL PolyJet transfection reagent (Fisher Scientific 50-478-8) was mixed with 250 µL DMEM. The DNA and PolyJet mixtures were then combined at a 1:1 ratio and incubated at room temperature for 30 minutes. The resulting transfection mixture was added dropwise to the cells and incubated for 16 hours. Media was then replaced with complete DMEM (10% FBS) supplemented with 5 µg/mL Blasticidin S HCl (Thermo Fisher Scientific A1113903) for selection. Cells were maintained under blasticidin selection for all subsequent experiments.

### Mitophagy measurements using mtKeima

HeLa cells expressing human MARCHF5 with G57 (human WT, squid recoded equivalent) or E57 (squid WT equivalent) under the control of a doxycycline-inducible promoter were seeded into live-cell imaging chambers (ibidi, 80806) at a density of 5x10^3^ cells/well. Upon reaching 70% confluence, the cells were induced with doxycycline (1 µg/mL), and transfected with the mtKeima construct^74^ provided by Ken Nakamura^75^ using Lipofectamine 2000 (Thermo Fisher Scientific, 11668019), following the manufacturer’s instructions. The cells were left for two days before mitophagy measurements. Culture medium was added alongside compounds for treatment, including either 0.1% DMSO or 10 µM FCCP (Sigma-Aldrich, SML2959). The cells were incubated for 1 hour prior to imaging. The cells were transferred to a confocal laser scanning microscope (Zeiss LSM880) fitted with a Plan-Apochromat 63x/1.4 Oil DIC M27 objective and PMT detector, with its incubation chamber pre-heated to 37°C and atmosphere filled with 5% CO2. Neutral mtKeima was imaged by excitation at 488 nm and detection between 572-710 nm, while simultaneously capturing transmitted light. Acidic mtKeima was imaged by excitation at 561nm and detection between 572-722 nm. The pinhole was set to 0.75 AU and 0.87 AU, respectively, and pixel scaling was set to 70 nm to capture mitochondrial structures. Five images were captured for each condition per replicate. Image analysis was performed in Fiji, and graphs were prepared in GraphPad Prism. A region of interest was generated of transfected cells based on the transmitted light, delimiting the area to be analyzed. A mask was generated of mitochondria by using both mtKeima channels. A minimum filter was first applied, followed by a maximum filter (radius=3). The Li AutoThreshold function was used to create a binary mask of mitochondria, followed by a dilation of the mask using EDM Binary Operations (iterations=3), watershed, and erosion using EDM Binary Operations (iterations=3). The resulting mask was used to create a region of interest to be analyzed. The region of interest, including all mitochondria, was split into individual pieces, and the intensity of each mitochondrion was measured from the raw, unprocessed image. The ratio of acidic to neutral mtKeima was calculated per mitochondrion. The results shown were obtained from three separate biological and technical replicate experiments.

### Seahorse analysis of mitochondrial respiration

Mitochondrial respiration was measured using the Seahorse XF Cell Mito Stress Test Kit (Agilent, 103015-100) following the manufacturer’s instructions. HeLa cells expressing human MARCHF5 with G57 (human WT, squid recoded equivalent) or E57 (squid WT equivalent) under the control of a doxycycline-inducible promoter were seeded into 96-well plates for Seahorse analysis (Agilent, 80806) at a density of 5x10^3^ cells/well. The next day, cells were induced with doxycycline (1 µg/mL) for two days before respiration analysis. The day before analysis, the sensor cartridge was hydrated in Seahorse XF Calibrant overnight at room temperature in a non-CO_2_ incubator. On the day of analysis, Seahorse XF DMEM (1 mM pyruvate, 2 mM glutamine, 10 mM glucose) was prepared and warmed. Cell culture medium was replaced, and the cells were placed into a non-CO_2_ incubator for 1 hour. During this time, fresh compounds were reconstituted fresh and loaded into the injection ports of the sensor cartridge, to yield the following final well concentrations upon injection: 2.5 µM oligomycin, 1 µM FCCP, 0.5 µM rotenone/antimycin A. After 1 hour had passed, the sensor cartridge was transferred to the cell culture microplate, and the plate was inserted into a Seahorse XFe96 analyser (Agilent). Following calibration and equilibration, the baseline respiration was measured by three cycles of 3 minutes mixing, 3 minutes measurement. This was followed by three sets of compound injections: First, oligomycin, then FCCP, and finally rotenone/antimycin A. Compound injections were performed, followed by four cycles of 4 minutes mixing and 4 minutes measurement. Respiration data were normalized to the initial baseline OCR as a proxy for cell density, and plotted in GraphPad Prism for visualization.

## Supplemental Figures

**Figure S1.**
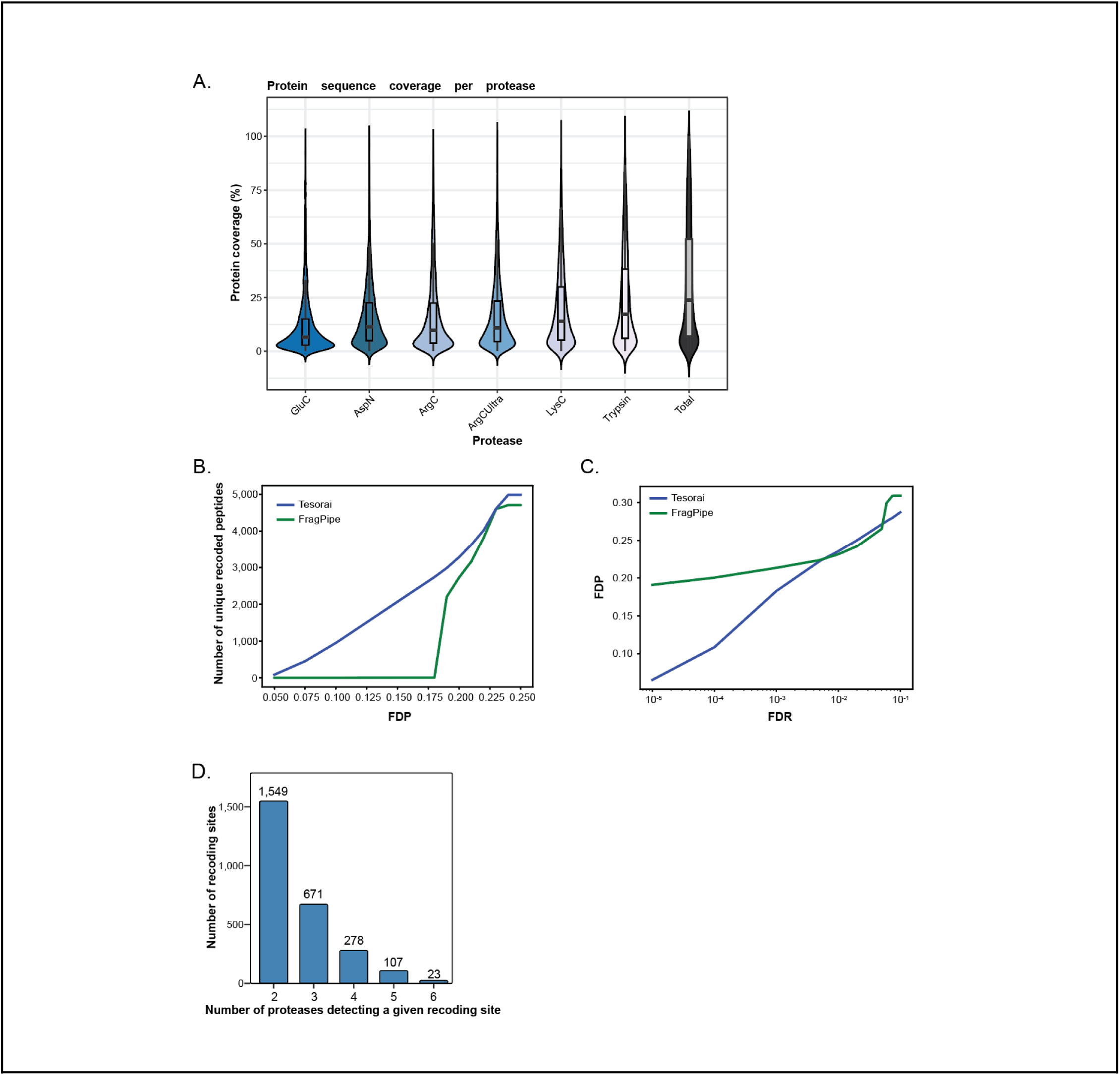
**A.** Violin plots for protein coverage provided by each protease, and total coverage for proteins identified in Fig. 1B. Comparison of Tesorai and FragPipe searches for FDP against recoded protein IDs (**B**) and FDR (**C**). **D.** Bar graph of the number of recoding sites identified across multiple proteases.

**Figure S2.**
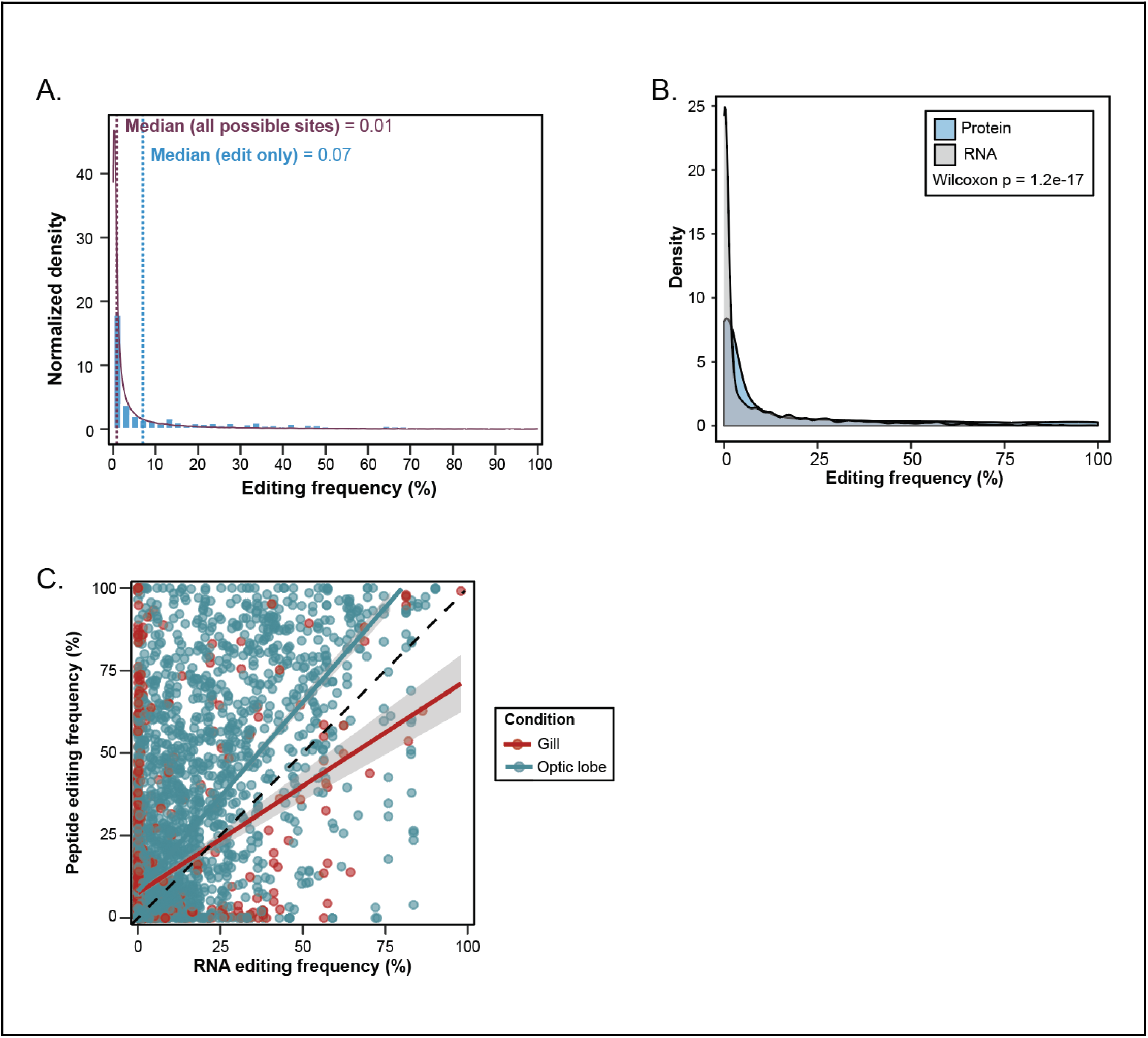
**A.** Distribution of editing frequencies between observed recoding sites and matching RNAseq sites, Wilcoxon-Cox p-value = 1.2e-17. **B.** Transcriptomic editing frequency for all possible editing sites compared to sites where only the recoded site was observed in the proteomics. **C.** Scatter plot of RNA editing frequency against peptide recoding frequency for all individual editing sites identified. RNA editing frequency is defined as the proportion of inosine-containing reads at a given site (I / [A + I]), while peptide editing frequency is defined as the proportion of recoded peptide signal relative to total peptide signal (recoded peptide intensity / [WT peptide + recoded peptide intensity]).

**Figure S3.**
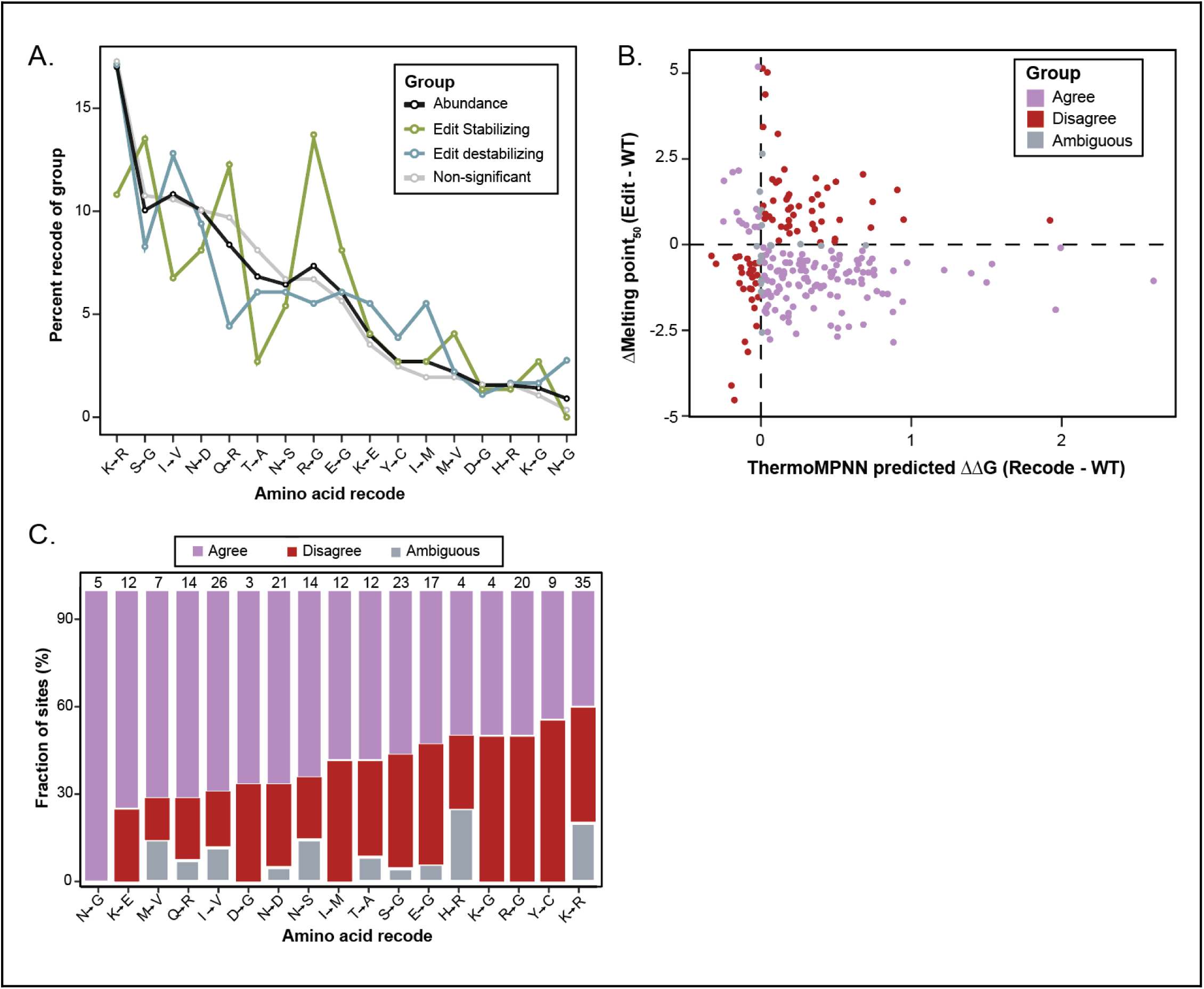
**A.** Distribution of single-site recoding events observed in Fig. 3B compared to sites seen from protein abundance experiments in Fig. 1A. **B.** Scatter plot for single-site recoding events from observed solvent shift Tm_50_ values and ThermoMPNN changes in free energy values. **C.** Bar plot for the fraction of recoded sites in agreement, disagreement, or that are ambiguous, broken down by amino acid recoding event. Each recoding type is normalized to 100%; n numbers are displayed above each bar.

**Figure S4.**
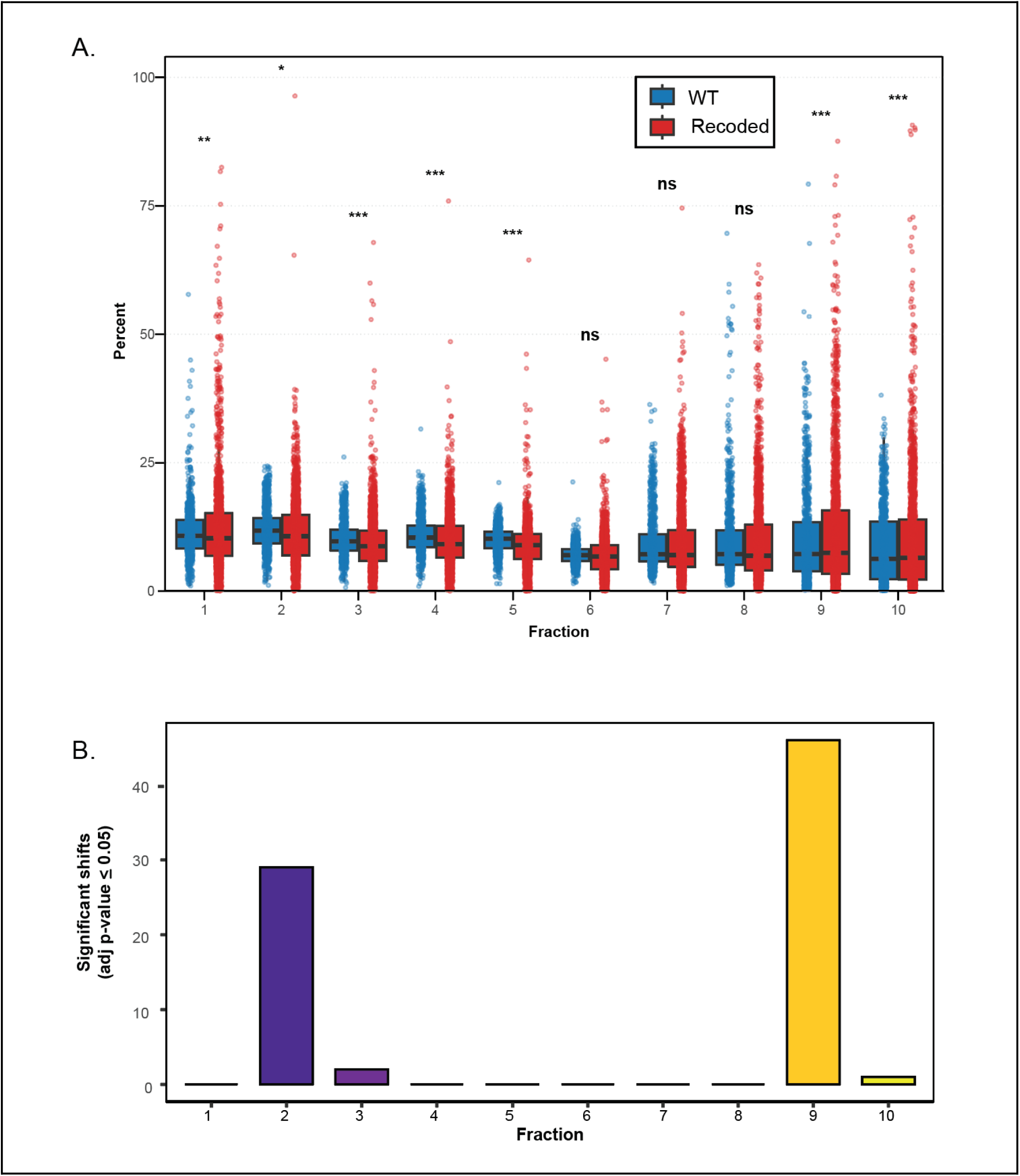
**A.** Box plot comparing the relative abundance of WT and matching recoded proteins across LOPIT-DC fractions. Significance markers based on BH corrected t-test, p-value < 0.05*, <0.01**, <0.001***. **B.** Number of significant shifts for a given comparison between the WT and recoded protein. Significance determined by BH corrected t-test, all entries p-value < 0.05.

**Figure S5.**
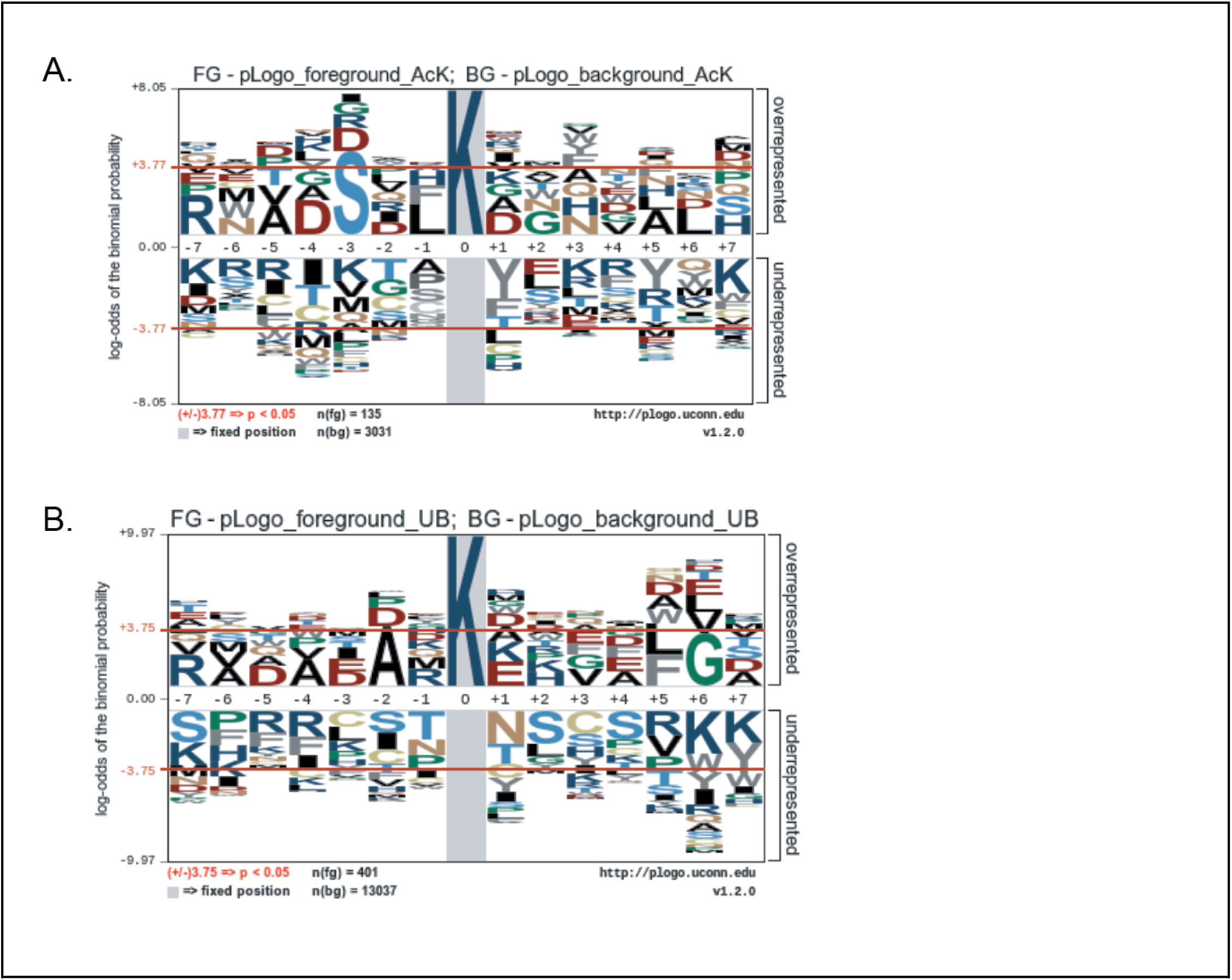
**pLogo motifs for (A) acetylation-ablative sites and (B) ubiquitination-ablative sites.**

**Figure S6.**
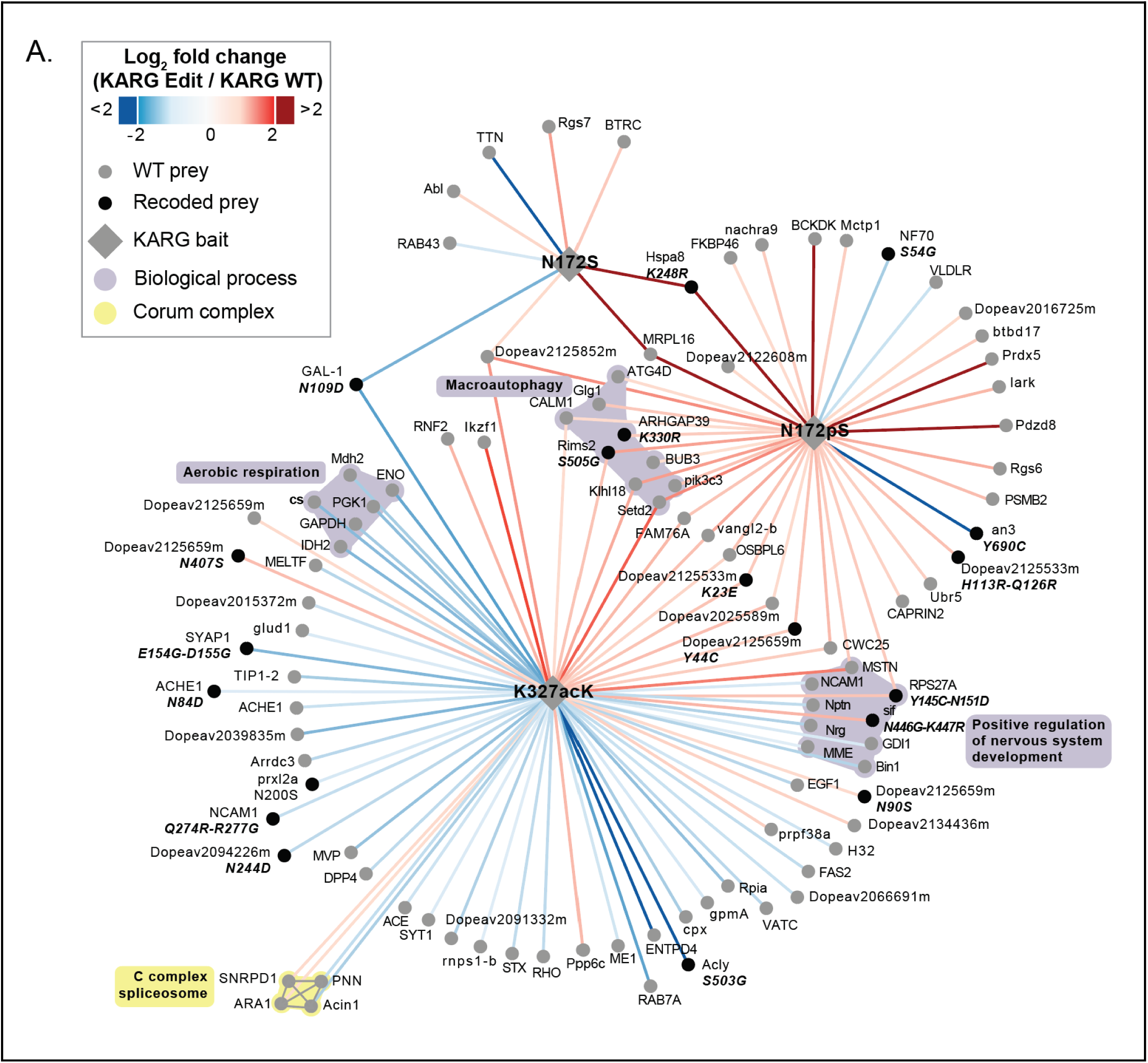
**A.** PPI network diagram for KARG K327acK, N172S, and N172pS sites. Biological processes and CORUM complexes are colored purple and yellow, respectively. Recoded-specific interactions are denoted in bold italics, with “-” between recoding events indicating a multi-recoded site. Sites were selected based on a Log_2_FC greater than or equal to |1|, adjusted p-value < 0.05, and CV less than 30% across replicates.

**Figure S7.**
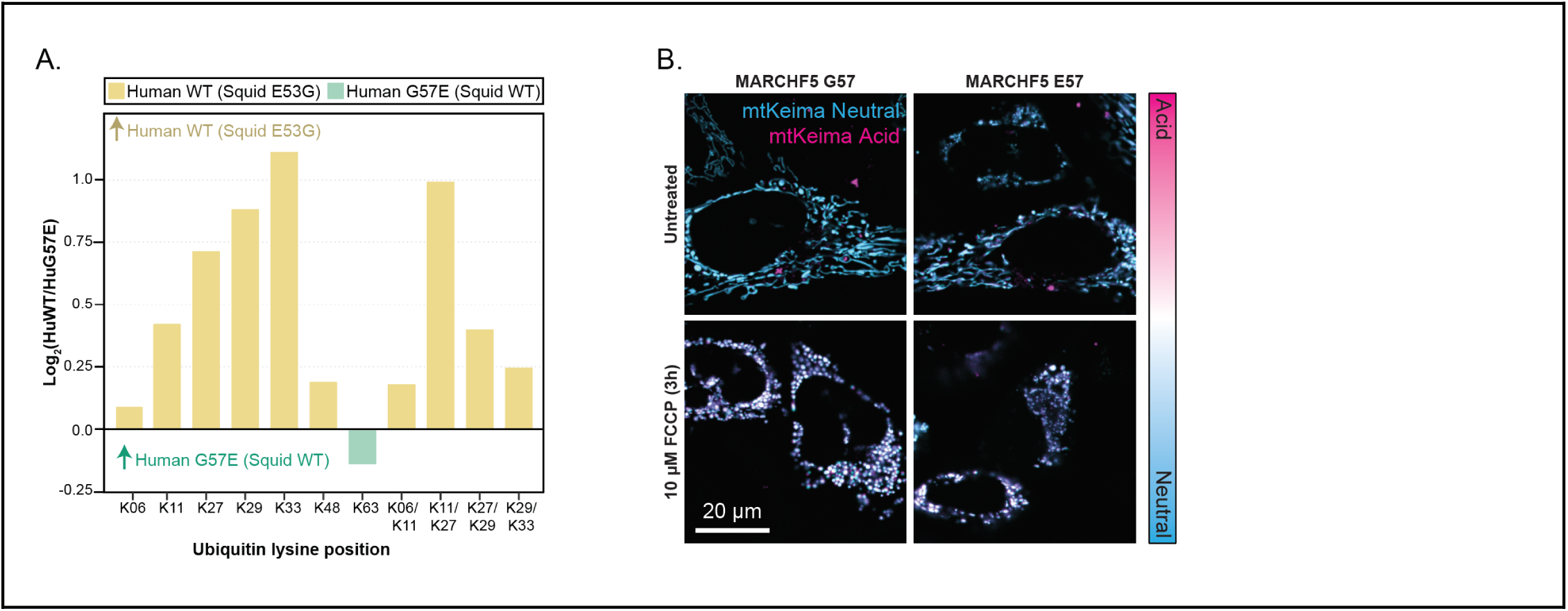
**A.** Log_2_FC for ubiquitin chain linkage positions between WT and G57E cells. **B.** Representative confocal microscopy image for mtKeima assay from Fig. 8G.

